# A conserved oscillatory system that positions the divisome in Archaea

**DOI:** 10.64898/2026.05.19.726155

**Authors:** Wei He, Ryusei Yoshida, Phillip Nußbaum, Kartikeyan Premrajka, Shamphavi Sivabalasarma, Chris van der Does, Marc Bramkamp, Sonja-Verena Albers

**Author notes:** Correspondence should be addressed to Sonja-Verena Albers. These authors contributed equally (Wei He, Ryusei Yoshida).

## Abstract

Precise placement of the cell division machinery is essential for cell division in most organisms, yet the mechanisms responsible for this process vary substantially across the domains of life. In Eukaryotes, cell division is primarily driven by ESCRT-based systems, while many Bacteria rely on the Min system to ensure accurate positioning of the division septum. By contrast, the mechanisms by which Archaea spatially regulate divisome assembly remain largely unknown. Here, we identify a three-protein system, which we term divisome positioning proteins A, B and C (DipA, DipB and DipC) that is essential for correct divisome positioning in Haloferax volcanii. Deletion of any dip gene results in mislocalization of cell division proteins and the formation of abundant minicells, similar to Min defects in bacteria. Furthermore, we show that all three Dip proteins undergo pole-to-pole oscillation, and that DipB and DipC assemble into ring-like structures at midcell. Biochemical analyses demonstrate that DipA is a membrane binding GTPase whose association with the membrane is disrupted by DipB. Notably, Dip homologues are widely conserved across diverse archaeal lineages that employ FtsZ-based division, indicating that the Dip system is a broadly distributed key regulator of FtsZ-based cell division in Archaea. Despite its striking functional similarities to the bacterial Min system, the Dip system is entirely unrelated at the sequence level, representing a compelling example of convergent evolution of oscillatory spatial regulators in distinct domains of life.

## Introduction

Cell division is a fundamental process for all living organisms but the mechanisms involved in this process vary considerably across all three domains of life. For example, in Eukaryotes, division is mainly driven by the ESCRT-based (Endosomal Sorting Complex Required for Transport) system^1^, whereas most bacteria employ the tubulin homolog FtsZ (filamentous temperature-sensitive gene Z)2. In Archaea, both systems are used, and while the ESCRT-based system evolved within the TACK Archaea, most Archaea use the FtsZ-based system^3–10^. Furthermore, in contrast to bacteria, most archaeal lineages carry two FtsZ homologues (FtsZ1 and FtsZ2)^5^. In *Haloferax volcanii*, the two FtsZ homologs have distinct functions: FtsZ1 scaffolds the Z-ring independent of FtsZ2 and provides a fundamental role in recruiting other division proteins and stabilizing the division machinery, whereas FtsZ2 seems to function primarily in cell constriction^5^. In addition to FtsZ1 and FtsZ2, an archaeal homologue of the bacterial SepF^6^, two PRC barrel domain proteins (CdpB1/2)^8,9^, and a recently reported transmembrane protein CdpA^10^ were also found to be part of the cell division machinery (the divisome) in *H. volcanii*.

In many bacteria, the precise placement of the divisome to midcell is dependent on the Min system, a negative regulatory system that was first identified through mutations that caused cell division defects and resulted in the production of minicells^11–13^. The Min system in the model bacterium *Escherichia coli* consists of three proteins: MinC, MinD and MinE^12^. Briefly, the MinC protein interacts with FtsZ, inhibiting its polymerization and preventing Z-ring assembly^14^. MinD is a membrane-associated ATPase, where it dimerizes and directs MinC to plasma membrane in the presence of ATP^15,16^. The membrane-bound MinD not only recruits MinC but also MinE, which displaces MinC from MinD and stimulates the MinD ATPase activity causing membrane detachment of MinD^16–18^. The MinD protein therefore re-binds ATP and attaches to the membrane on the opposing cell pole, where it subsequently recruits MinC and MinE, ultimately establishing a pole-to-pole oscillatory cycle^19–22^. The oscillation of the Min system between cell poles results in a net minimum concentration of MinCD at midcell, thus allowing FtsZ polymerization and Z ring assembly to occur near midcell^12,14,23^.

While the positioning of the divisome in bacteria has been well characterized, the current knowledge about the spatiotemporal coordination of archaeal cell division is much more limited. Previous studies examining the division plane location in *H. volcanii* and *Haloarcula japonica* have suggested a MinCDE-like mechanism for division site placement in these Archaea^24^. A few members of Euryarchaeota were indeed found to encode MinD homologs, although other Min components (i.e., MinC and MinE) are absent^7^. However, detailed analyses of the four MinD homologs in *H. volcanii* showed that none of these homologs is directly involved in cell division^25^. Instead, MinD2 and MinD4 modulate the positioning of the chemotaxis arrays and archaellum motors, thereby affecting cell motility^25–27^. Therefore, the systems responsible for the precise positioning of the archaeal divisome remain unknown. Here, we show that an oscillatory system unrelated to Min, consisting of *hvo_3012, hvo_3013* and *hvo_3014*, is widely conserved across Archaea and regulates the positioning the archaeal division machinery at midcell. Consequently, we name it as the Dip (Divisome positioning) system.

## Results

### The Dip system is widely distributed in Archaea

We have previously shown, that SepF is essential for cell division in H.volcanii^6^. Interestingly, HVO_3013 is structurally highly homologous to SepF (HVO_0392), although HVO_3013 seems to lack a membrane targeting sequence like SepF^28^ (Supplementary Figure 1a). A previous study showed that *hvo_3012* (*dipC*), *hvo_3013* (*dipB*) and *hvo_3014* (*dipA)* form an operon near the orc1 (origin recognition complex) gene in *H. volcanii*, with a strong primary transcription start site (TSS) upstream of *hvo_3014* and a minor TSS in the 3’-region of hvo_3014^29^ (Fig. 1a).

**Fig. 1:**
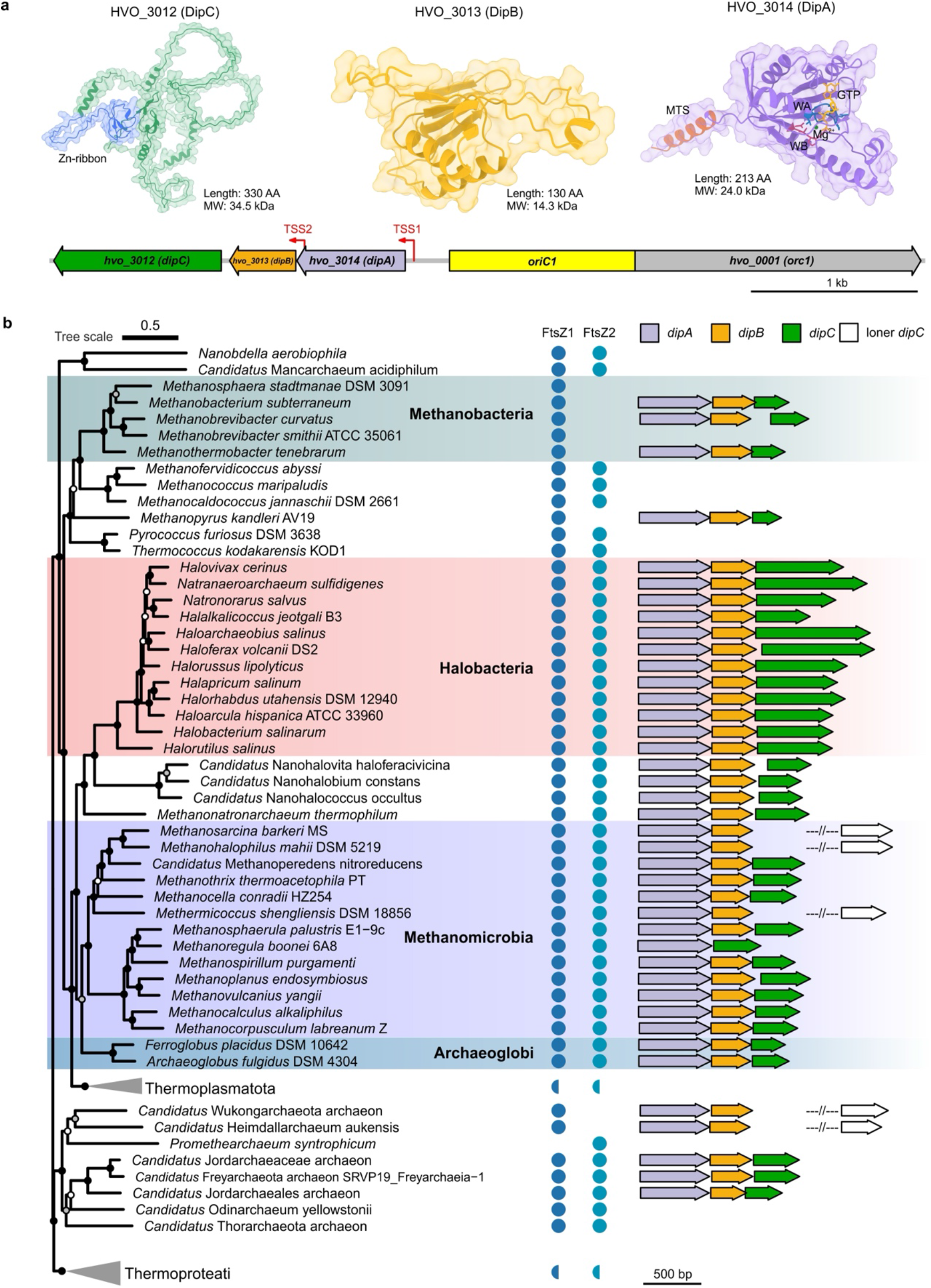
The Dip system is widely conserved in Archaea. **a**, Upper: Alphafold 3 predicted structures of HVO_3012, HVO_3013 and HVO_3014. The Zn-ribbon domain of DipC (blue), membrane targeting sequence (MTS, chocolate) of DipA, Walker A motif (WA, blue) and Walker B motif (WB, red) of DipA, the GTP (orange) and Mg^2+^ (green) binding to DipA are colored separately. Protein sequence length and molecular weight (MW) as indicated. Lower: genomic organization of *dip* genes, replication origin (*oriC1*) and *hvo_001* (*orc1*) in *H. volcanii*. Two transcription start sites (TSS) are indicated by red arrow. Scale bar, 1 kb. **b**, The phylogenetic tree of Archaea (re-rooted at midpoint) with the presence of FtsZ1, FtsZ2 homologs and dip operon. Semicircles indicate that the corresponding protein couldn’t be identified in all the taxa of this clade. The loner *dipC* gene outside the operon is indicated by unfilled arrow and double slashes. The local support values (LSV) based on the Shimodaira-Hasegawa (SH) test are indicated by dark dots (0.9 < LSV ≤ 1), gray dots (0.7 < LSV ≤ 0.9) and white dots (LSV ≤ 0.7). Tree scale as indicated. Scale bar in genomic organization, 500 bp.

DipC is the largest Dip protein in *H. volcanii* (330 amino acids, 34.5 kDa) and contains an N-terminal Zn-ribbon domain (Fig. 1a). AlphaFold prediction indicates that its serine-rich central region is highly disordered^30^. Residues 32-270 were predicted by FuzDrop^31^ as a droplet-promoting region (DPR) with high droplet-promoting probabilities (pDP), suggesting a strong propensity for spontaneous liquid-liquid phase separation and condensate formation (Supplementary Figure 1b). DipB (130 amino acids, 14.3 kDa) contains a DUF2073 domain of unknown function (Fig. 1a). DipA is a predicted GTP-binding protein with conserved GTP-binding motifs and has been proposed to drive extracellular vesicle formation in haloarchaea^28^. The Walker A motif (residues 31-38, GPPNAGKT) forms a loop between a β-strand and an α-helix, while the Walker B motif (residues 87-90, DTPG) is located downstream at the end of a β-strand (Fig. 1a and Supplementary Figure 1c). Its N-terminus contains a membrane-targeting sequence predicted to form an amphipathic helix enabling membrane interaction (Supplementary Figure 1d).

To determine how conserved the Dip system is in the Archaea, we analyzed 766 archaeal genomes from NCBI database using MacSyFinder v2^32^(Supplementary Data 1). The Dip system was found in 559 of 650 Methanobacteriati (formerly “Euryarchaeota”) genomes, including the classes Methanobacteria, Halobacteria, Methanomicrobia and Archaeoglobi, but was not present in Methanococci and Thermococci (Supplementary Data 2 and Supplementary Figure 2). Despite its wide distribution in Methanobacteria and Methanomicrobia, the Dip system was not found in genomes of *Methanosphaera, Methanogenium* and some species of *Methanobrevibacter*. Notably, the dipB gene in Methanoregula was missing (Supplementary Figure 2). Besides Methanobacteriati, the Dip system was also present in some species of the Promethearchaeati (formerly “Asgard”) and Candidatus Nanohalarchaeota within Nanobdellati (formerly “DPANN superphylum”) (Supplementary Figure 2). Intriguingly, the Dip system was only present in Archaea employing the FtsZ-based cell division system, although its presence did not correlate with FtsZ2 (e.g., Methanobacteria) (Fig. 1b, Supplementary Figure 2, and Supplementary Data 3), suggesting a functional dependence on FtsZ1.

While DipA and DipB homologs are highly conserved across their entire sequences, DipC homologs show conservation mainly at their N- and C-terminal regions^29^, whereas the central region is variable (Supplementary Figure 3). The observed length differences are primarily due to variation in the intrinsically disordered region (IDR) located between the conserved termini. Accordingly, DipC homologs in Halobacteria (276.021 ± 60.786 amino acids, mean ± SD, n = 416) are substantially longer than other homologs (131.526 ± 19.584 amino acids, mean ± SD, n = 152) (Supplementary Figure 3 and Supplementary Data 2). In addition to the large variation in DipC length, the distance between *dipC* and *dipAB* also varied among species (Fig. 1b). In the family *Methanosarcinaceae, Methermicoccus shengliensis* DSM 18856, *Methanobrevibacter filiformis* and some Asgard Archaea, the *dipC* gene does not cluster with *dipAB*, hence it is labelled as loner gene (inter genes > To explore the function of the Dip proteins, we first 3) in this study (Fig. 1b and Supplementary Figure 2). constructed single gene deletion strains of *dipA, dipB* and *dipC* in *H. volcanii* H26. When comparing the **Deletion of *dip* genes affects cell division and** growth curves of knockout strains with wild-type H26 **produces minicells** strains, Δ*dipA* and Δ*dipC* had slightly decreased growth rate, while the growth rate of Δ*dipB* was unaffected (Supplementary Figure 4a, b). Consistent with this, Δ*dipA* and Δ*dipC* cells showed slightly reduced viability compared with H26 (Supplementary Figure 4c). Although growth was not strongly affected, the Δ*dipA* and Δ*dipC* strains produced a large amount of minicells which sometimes could not proliferate (Fig. 2a, c, d and Supplementary Movie 1). Similarly, Δ*dipB* also produced minicells, but it also led to more elongated and enlarged cells, resulting in a larger cell area overall (Fig. 2b, d). The minicells from all three knockout strains could either contain DNA or be DNA free (Supplementary Figure 4d). The wild-type H26 cells normally constrict at the middle of cells and produce two daughter cells with similar size and shape (Supplementary Movie 1). By contrast, the Dip mutants could constrict and divide simultaneously at multiple sites. The placement of those division sites in the mutants lacked an obvious pattern, with some sites close to the cell poles and others along the cell edges (Fig. 2a-c and Supplementary Movie 1). Surprisingly, the deletion of dip genes also caused severe motility defects, with Δ*dipA* and Δ*dipC* cells being completely non-motile (Supplementary Figure 4e, f).

**Fig. 2:**
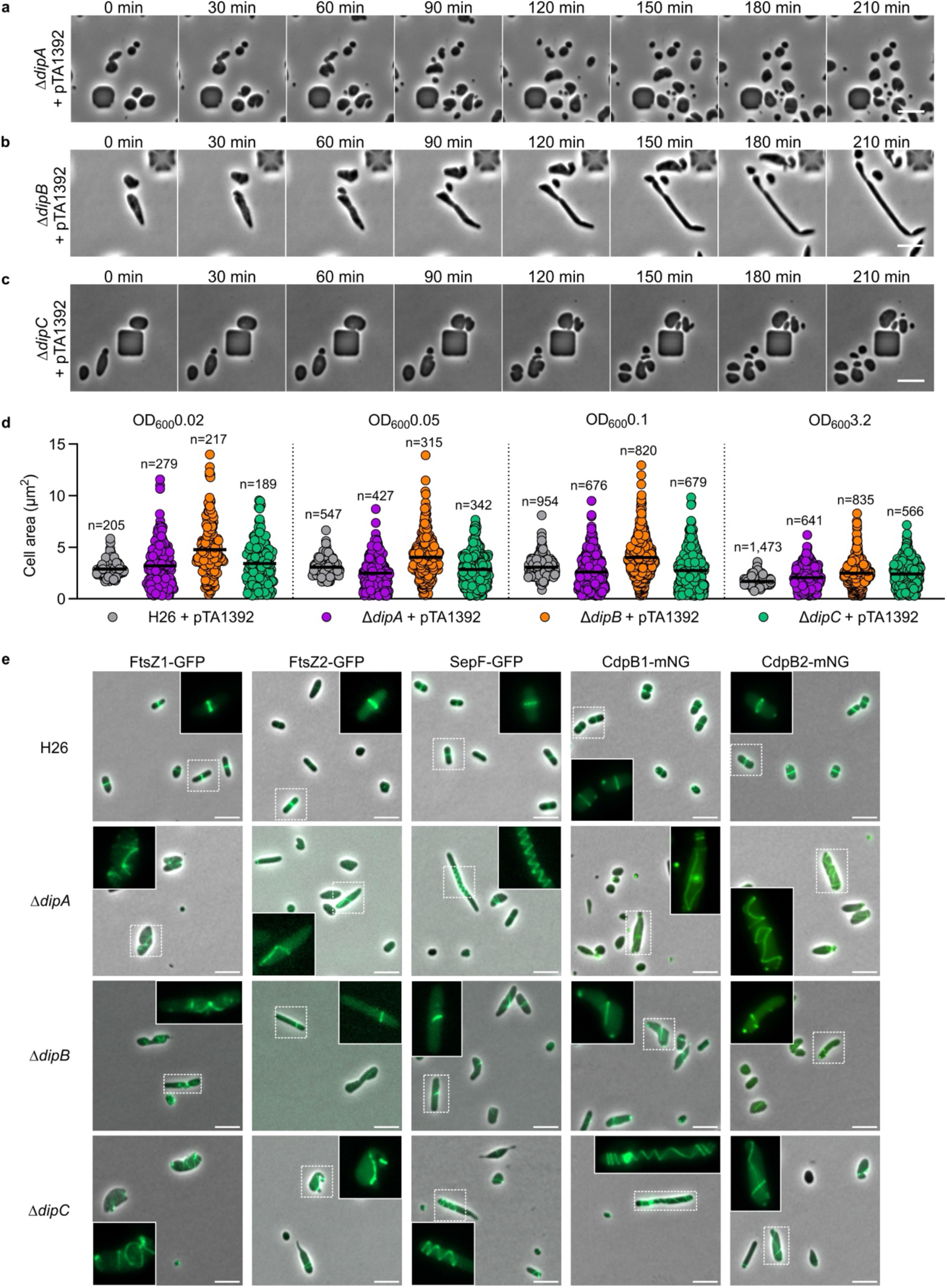
Deletion of *dip* genes affects cell division. Time-lapse microscopy of (**a**) Δ*dipA*, (**b**) Δ*dipB*, and (**c**) Δ*dipC* with empty plasmid (pTA1392) in microfluidic chambers. Only parts of the complete videos are shown. **d**, Cell area analysis of H26, Δ*dipA*, Δ*dipB* and Δ*dipC* during different growth stage (OD_600_, 0.02 early exponential phase, 0.05-0.1 exponential phase, 3.2 stationary phase), mean values are shown as dark lines. All strains were transformed with empty expression plasmid pTA1392, data were taken from 3 independent replicates. **e**, Fluorescence microscopy of wild-type strain (H26), Δ*dipA*, Δ*dipB*, and Δ*dipC* expressing FtsZ1-GFP, FtsZ2-GFP, SepF-GFP, CdpB1-mNG (mNeonGreen) and CdpB2-mNG. Fluorescent channel image of cells in dashed boxes are magnified. All cells were grown in liquid Hv-Cab medium. All scale bars, 5 µm.

**Fig. 2:**
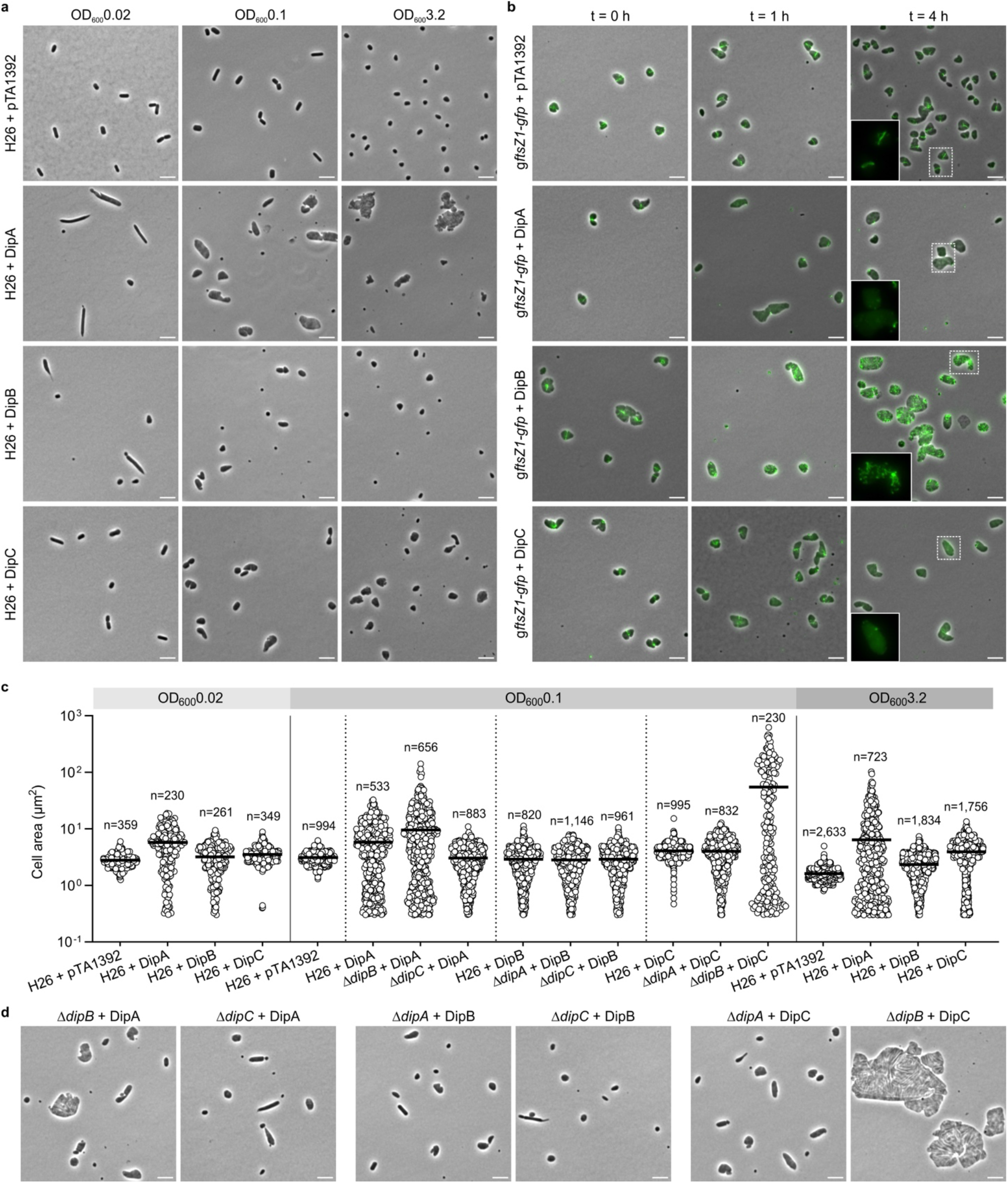
Overexpression of Dip proteins impacts cell division and divisome localization. Phase-contrast images (**a**) and cell area analysis (**c**) of *H. volcanii* wild-type H26 cells with empty vector (pTA1392) and Dip protein overexpression plasmids (DipA, pSVA6849; DipB, pSVA6848; DipC, pSVA6847) from different growth stage in liquid Hv-Cab medium + 1 mM tryptophan. **b**, Fluorescence microscopy of *H. volcanii* HTQ971 (*gftsZ1-gfp*, with genomic integration of a *gfp* tag to *ftsZ1* C-terminal) cells with empty vector (pTA1392) and Dip protein overexpression plasmids. Cells were imaged before induction (t = 0) and different time points (1 and 4 h) after induction. The expression of Dip proteins was induced by adding 1 mM tryptophan. Fluorescent channel image of cells in dashed boxes are magnified at lower left. Phase-contrast microscopy (**d**) and cell area analysis (**c**) of the indicated strains (Δ*dipA*, Δ*dipB* and Δ*dipC*) expressing the Dip proteins (DipA, DipB and DipC) or containing empty vector at exponential phase (OD_600_ about 0.1) in Hv-Cab medium + 1 mM tryptophan. The area data shown merged all 3 datasets. Mean values are shown as dark lines in (**c**). All scale bar, 5 µm.

We then characterized the localization of divisome proteins in Dip mutants compared with wild-type H26 cells. Fusion proteins (using mNeonGreen (mNG) or GFP) of FtsZ1, FtsZ2, SepF, CdpB1 and CdpB2 localized at midcell and formed single ring-like structures in wild-type H26 cells (Fig. 2e and Supplementary Movie 2). By contrast, these cell division fusion proteins were randomly distributed in the dip deletion mutants (Fig. 2e and Supplementary Movie 2). Nevertheless, FtsZ1 and FtsZ2 co-localized in both the wild-type and knockout strains (Supplementary Figure 5). In Δ*dipA* and Δ*dipC* cells, FtsZ1 could polymerize into more filaments, thereby probably promoting the assembly of other cell division proteins into filamentous structures (Fig. 2e and Supplementary Figure 5b, d). However, the localization of those filaments was not restricted and these proteins sometimes formed spiral-like structures that could be observed throughout the whole cell (Supplementary Figure 5b, d). By contrast, the FtsZ1 filaments in Δ*dipB* cells were much shorter, with a portion of FtsZ1 aggregating into clusters (Fig. 2e and Supplementary Figure 5c). Consequently, the divisome failed to assemble into complete ring in some ΔdipB cells (Fig. 2e).

Using 3D structured-illumination microscopy (3D-SIM), we found that FtsZ1 could form clear flattened ring-like structure in wild-type H26 cell, which is consistent with a previous study^5^ (Supplementary Figure 6a and Supplementary Movie 3). 3D-SIM analysis showed that the localization of FtsZ1 filaments in Δ*dipA* cells was closer to the cell membrane, while it was more cytoplasmic in ΔdipC cells (Supplementary Movie 3). FtsZ1 in Δ*dipB* cells, however, formed less filaments (Supplementary Movie 3).

To further investigate the effect of Dip proteins on FtsZ1 dynamics and localization, we performed single-particle tracking (SPT) by fusing FtsZ1 with HaloTag. The single molecule dynamics showed that FtsZ1 was more mobile in Dip protein knockout strains than in the wild-type H26 strain (Supplementary Figure 6b). The distribution of the molecule tracks was also different among H26, Δ*dipA*, Δ*dipB*, and Δ*dipC* cells. In H26 cells, most tracks appeared at midcell, but the tracks were more near to the membrane in Δ*dipA* cells, consistent with the 3D-SIM results (Supplementary Figure 6c, d). In contrast, the FtsZ1 tracks were found all over the cytoplasm in absence of DipC, which is also in line with the data obtained by 3D-SIM (Supplementary Figure 6c, d).

### Overexpression of Dip proteins impacts cell shape and the localization of cell division proteins

We then overexpressed Dip proteins under a tryptophane-inducible promoter in wild-type and dip knockout strains to assess whether they have distinct or overlapping functions. Overexpression of DipA in H26 cells caused cell elongation during early exponential phase and swelling during exponential and stationary phases, accompanied by formation of numerous minicells and small cell debris (Fig. 3a, c). This abnormal morphology suggests that biomass production continued while cell division was impaired, leading to ongoing cell expansion. We also overexpressed DipA in strains carrying genomic ftsZ1-gfp or sepF-gfp fusions. Although the GFP tag on ftsZ1 caused mild division defects, FtsZ1-GFP still formed rings at midcell, positioned away from the cell pole in H26 cells (Fig. 3b). By contrast, overproduction of DipA resulted in FtsZ1-GFP foci rather than ring structures (Fig. 3b). The genomic integration of the gfp tag to the sepF locus resulted in long filamentous cells but did not affect the ring formation of SepF-GFP (Supplementary Figure 7a). However, when overexpressing DipA in these filamentous cells, SepF-GFP rings became widened and dispersed, and in some cells, SepF-GFP stayed in one of the cell poles as foci or patches (Supplementary Figure 7a).

**Fig. 3:**
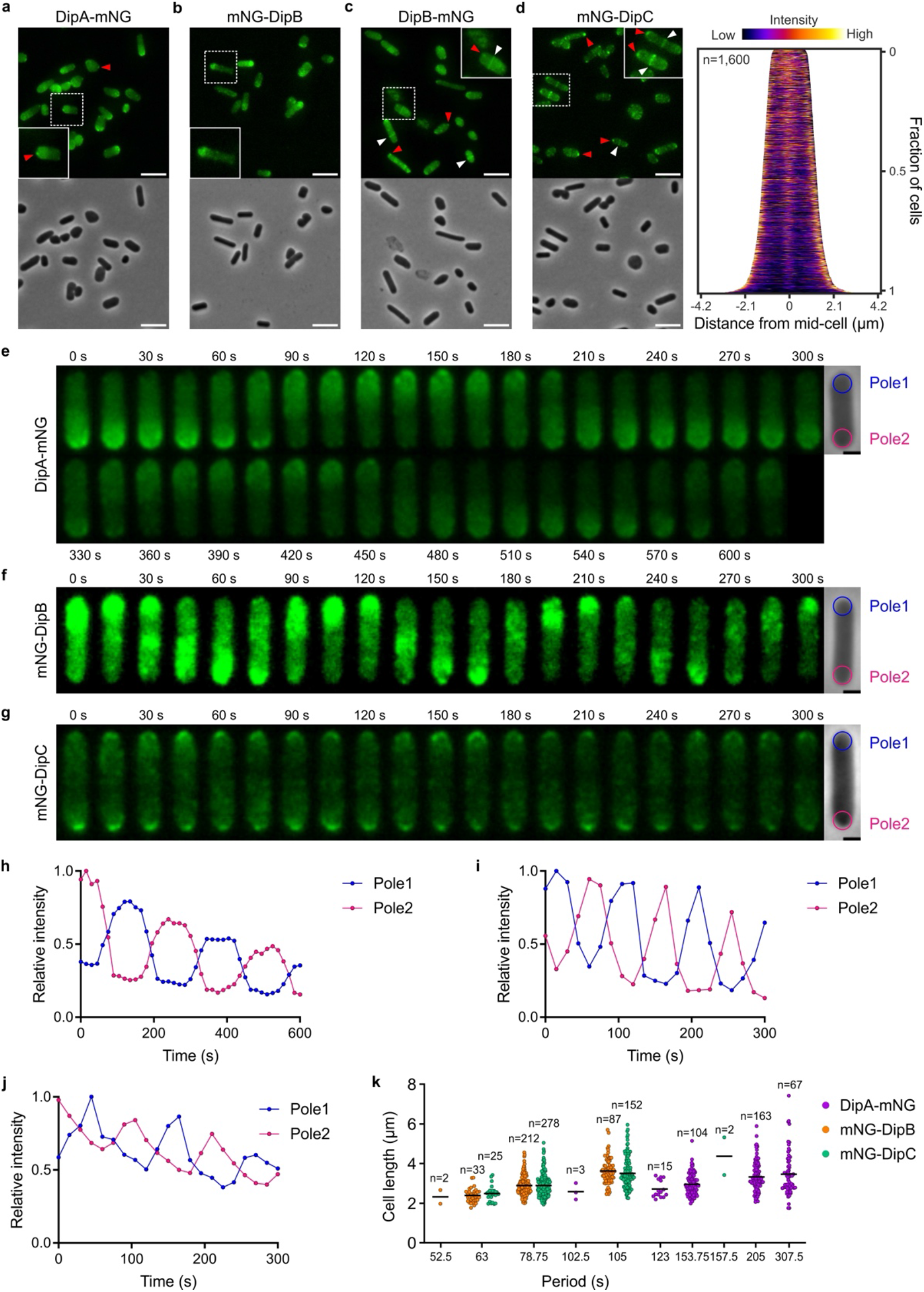
Dip is an oscillatory system. Fluorescence microscopy of *H. volcanii* H26 cells in the exponential phase expressing (**a**) DipA-mNG, (**b**) mNG-DipB, (**c**) DipB-mNG and (**d**) DipC-mNG under control of the tryptophan promotor. Cells were grown in liquid Hv-Cab medium and induced with 1 mM tryptophan for 4 h. Demographic analysis in (**d**) shows signal intensity of DipC in cells, data were taken from 3 independent replicates. Fluorescent channel images of representative cells in dashed boxes are magnified. Representative foci are indicated by red arrowheads, ring-like structures are indicated by white arrowheads. Kymograph of (**e**) DipA-mNG, (**f**) mNG-DipB and (**g**) mNG-DipC oscillation in H26. Images were taken every 15 s for 5 min or 10 min. Two pole areas (diameter, 1 µm) are labelled, the average fluorescence intensity in those areas were calculated by ImageJ and plotted (see below). Relative fluorescence intensity of two cell pole areas (indicated in **e-g**) of (**h**) DipA-mNG, (**i**) mNG-DipB, and (**j**) mNG-DipC. **k**, The distribution of oscillation periods (based on the intensity ratio of two cell poles) observed for each mNG construct and cell length, data were taken from 3 independent replicates, mean values are shown as dark lines. Scale bar in (**a-d)**, 5 µm, (**e-g)**, 1 µm.

In contrast to DipA, overproduction of DipB in H26 cells did not result in enlarged cells, but led to the formation of a large number of minicells (Fig. 3a, c). Consistent with this, FtsZ1-GFP formed numerous filaments within the cells upon increased DipB expression, with the filaments formed at random places within the cells (Fig. 3b). Similarly, SepF-GFP could form up to 7 rings per cell when DipB was overexpressed, and the number of rings increased as the cell length increased (Supplementary Figure 7a, b). This indicated that the overexpression of DipB could remove the positional restriction of FtsZ1 polymerization while maintaining its ability to recruit downstream cell division proteins.

The overexpression of DipC in H26 cells also resulted in an increased cell volume, although this was less pronounced than the increase caused by DipA overexpression (Fig. 3a, c). FtsZ1-GFP started to aggregate into foci after DipC overexpression, and only a few FtsZ1-GFP filaments remained after 4 hours of induction of DipC overexpression (Fig. 3b). The dispersing and breakdown of division machinery could be observed in sepF-gfp cells after DipC overproduction (Supplementary Figure 7a). Interestingly, in DipC overexpression cells, SepF-GFP was not present at the cell poles like in DipA overexpression cells (Supplementary Figure 7a).

We then overexpressed Dip proteins in the various dip deletion strains to verify the functional interdependence of these proteins. The overexpression of DipA in Δ*dipB* cells showed similar effect as in H26, where cell division was heavily inhibited (Fig. 3c, d). However, DipA overexpression in Δ*dipC* cells did not lead to enlarged cells, but a large number of minicells (Fig. 3c, d). These results suggest that the inhibition of DipA on cell division is dependent on DipC. Surprisingly, overexpression of DipC in Δ*dipB* cells caused nearly completed division defects and produced giant cells (with the area reaching hundreds of µm2) (Fig. 3c, d). Finally, the overexpression of Dip proteins in H26 cells also significantly reduced motility (Supplementary Figure 8).

### Dip proteins oscillate and localize to the cell division site

To investigate the subcellular localization of the Dip proteins, we then expressed C-terminal mNG-tagged DipA proteins (DipA-mNG) from plasmids under the control of a tryptophan inducible promotor in wild-type H26 cells. DipA-mNG predominantly localized in one of the halves of the rod-shaped cells, with small foci present near the poles in some cells (Fig. 3a). Time-lapse imaging revealed that DipA-mNG proteins oscillated along the long axis of cells while the small foci kept stationary (Fig. 3e, h and **Error! Reference source not found**.). DipA-mNG showed an oscillation period of approximately 205 s in most cells, and the oscillation period increased along the cell length (Fig. 3k).

Similar to DipA-mNG, mNG-DipB also clearly localized at one of the cell halves and showed pole-to-pole oscillation in H26 cells (Fig. 3b, f and **Error! Reference source not found**.). However, mNG-DipB showed much shorter dwelling time at the poles and a shorter oscillation period of approximately 78.75 s in most cells, which increased along the cell length (Fig. 3i, k). The DipB-mNG could also oscillate, but showed a stronger diffused signal compared to the N-terminal fusion. DipB-mNG also formed small foci that were spatially and temporally static (Fig. 3c and **Error! Reference source not found**.). Intriguingly, DipB-mNG also formed stable ring-like structures in some cells and co-localized with FtsZ1 (Fig. 3c and **Error! Reference source not found**.a). These ring-like structures were not always at the cell centre, possibly due to overexpression of DipB-mNG (Fig. 3c).

DipC displayed several localization patterns, including pole-to-pole oscillation, formation of ring-like structures at midcell and polar foci (Fig. 3d, g and **Error! Reference source not found**.). mNG-DipC had a similar oscillation period to mNG-DipB (Fig. 3j, k). While mNG-DipC ring-like structures co-localized with FtsZ1 in some cells (**Error! Reference source not found**.b), these structures were highly dynamic and unstable, forming and disappearing multiple times during a single cell cycle (**Error! Reference source not found**.). Unlike the static foci formed by DipA-mNG and DipB-mNG, mNG-DipC resulted in bright foci or cap-like structures with changing intensity during the period of oscillation (Fig. 3g and **Error! Reference source not found**.).

### DipA and DipB form an independent oscillator that drives oscillation of DipC

As described above, the functional relationship between Dip proteins prompted us to further investigate whether their cellular localizations are interdependent. The pole-to-pole oscillation of DipA-mNG was not affected in the absence of DipC, while no DipA-mNG oscillation was observed in Δ*dipB* cells (Fig. 5a and Supplementary Movie 6). Consistently, the deletion of *dipA* also abolished the oscillation of DipB, whereas the absence of DipC had no effect on DipB localization (Fig. 5b, c and Supplementary Movie 6). Notably, despite the loss of oscillation of DipB in Δ*dipA* cells, the ring-like structure formed by DipB-mNG remained unaffected in the absence of DipA (Fig. 5b). The cellular position of mNG-DipC was fully dependent on DipA, as all localization patterns of mNG-DipC seen in H26 cells disappeared in Δ*dipA* cells (Fig. 5c and Supplementary Movie 6) By contrast, the deletion of *dipB* only disrupted the oscillation of mNG-DipC but did not affect its ring-like structure and polar focus (Fig. 5c and Supplementary Movie 6). Consistently, DipC exhibited higher dynamics in Δ*dipA* cells compared to H26 cells, but appeared more static in Δ*dipB* cells (Supplementary Figure 10). Taken together, DipA and DipB jointly drive the oscillation of the entire Dip system, whereas DipC appeared to act as a cargo. Moreover, the oscillatory proteins and the ring-forming proteins seem to represent two independent pools.

### DipA is a membrane binding GTPase

DipA was predicted to be a GTP-binding protein with a Walker A and a Walker B motif (Fig. 1a). To test whether DipA binds and hydrolyses GTP, we tried to whether DipA binds and hydrolyses GTP, we tried to purify *H. volcanii* DipA. Unfortunately, this protein was highly unstable *in vitro*. Therefore, we purified the DipA homolog from the hyperthermophilic archaeon *Archaeoglobus fulgidus* (AfDipA, Supplementary Figure 11a). The purified AfDipA was able to efficiently hydrolyze GTP at 80 °C, but not ATP or CTP (Fig. 5d and Supplementary Figure 12a, b).

To investigate the role of GTP hydrolysis on the function of DipA, we mutated the Walker A motif (K37A) and Walker B motif (D87A) of *H. volcanii* DipA, and checked the localization of the mutants (DipAK37A-mNG and DipAD87A-mNG) expressed from plasmids in H26 cells. Both DipAK37A-mNG and DipAD87A-mNG were not able to oscillate compared to DipA-mNG in H26, which indicated that the oscillation of DipA depends on the hydrolysis of GTP (Fig. 5e and Supplementary Movie 7). Interestingly, the expression of DipAK37A and DipAD87A in H26 cells also caused cell division defects and led to the production of minicells, suggesting that DipAK37A and DipAD87A interfere with native DipA in *H. volcanii* (Supplementary Figure 12c).

DipA has a predicted N-terminal membrane-targeting sequence (MTS), suggesting that it can associate with the cell membrane (Supplementary Figure 1d). To assess the association of purified AfDipA with phospholipids in vitro, we performed phospholipid vesicle sedimentation assay^17^. We found that 90% of AfDipA co-sedimented with the lipid vesicles (liposome) in the presence of GTP (Fig. 5f). In the presence of GDP, AfDipA did not increase in the pellet fraction, indicating that the GTP-bound form of AfDipA had a significantly higher affinity for phospholipids than the GDP-bound form (Fig. 5g). The binding of AfDipA to liposomes also significantly enhanced its GTP hydrolysis activity (Fig. 5f). By contrast, much less AfDipAL7R mutant (predict to break the hydrophobic face of MTS) was found in the pellet fraction after incubating with liposomes (Fig. 5f).

### DipB detaches DipA from the membrane

As described above, DipA and DipB are interdependent for oscillation in vivo. To investigate how they interact in vitro, the *A. fulgidus* DipB homologue (AfDipB) was also heterologously expressed and purified from *E. coli* (Supplementary Figure 11b and Supplementary Figure 12a). Interestingly, AfDipB eluted in SEC at a position corresponding to molecular weight about 27 kDa, corresponding to dimerized AfDipB. This dimer remained stable even under extreme high temperature condition, consistent with the hyperthermophilic growth condition of *A. fulgidus* (Fig. 5h). The purified AfDipB was incubated with AfDipA and liposomes at 80 °C for 30 min in the presence of GTP to determine the effect of AfDipB on AfDipA membrane association and GTP hydrolysis. Addition of AfDipB resulted in a significantly reduced (from ∼90% to ∼50%) amount of liposome-associated AfDipA (Fig. 5i).

To determine whether AfDipB detached AfDipA that was already bound to the liposome or directly sequestered AfDipA from binding to the liposome, we incubated AfDipA with liposome and GTP first, followed by addition of AfDipB to the reaction mixture (Supplementary Figure 12d). After incubating with AfDipB, the prebound AfDipA was removed from liposome and only 60% of AfDipA still co-sedimented with lipid vesicles (Supplementary Figure 12d). This supports the hypothesis that AfDipB detaches AfDipA from membranes rather than preventing AfDipA binding to the membrane. We also found that AfDipB slightly stimulated the GTPase activity of AfDipA in the absence of phospholipid vesicles (Supplementary Figure 12e). The stimulation was further enhanced when incubated together with phospholipid vesicles (Supplementary Figure 12e). Although DipA is a GTPase, the interactions that we observed in these experiments are reminiscent of the mechanism that has been shown in the Min System (MinC/D/E)^17^.

### DipB binds to membrane

In the Min system, not only MinD being able to associate with cell membrane, MinE can also directly interact with the membrane33. In the absence of AfDipA, 30% of AfDipB co-sedimented with liposomes, suggesting its membrane binding activity (Fig. 5i). Furthermore, when co-incubated with AfDipA, a larger fraction (additional 20%) of AfDipB co-sedimented with liposomes (although this increase in membrane affinity may be due to its interaction with AfDipA) (Fig. 5i). While DipB does not have a membrane targeting sequence like DipA or SepF6, it contains a positively charged loop (*H. volcanii*, amino acids: 85-99; *A. fulgidus*, amino acids: 73-85) with a small lipophilic face that may enable membrane binding (Supplementary Figure 1b and Supplementary Figure 11b). To investigate the function of this loop, we expressed and purified a DipB variant with a truncated loop region from *A. fulgidus* (AfDipBΔLoop). We then assessed the effect of AfDipBΔLoop on AfDipA membrane association and GTP hydrolysis by incubating AfDipBΔLoop and AfDipA in the presence of liposome and GTP. Similar to the native AfDipB, AfDipBΔLoop could also detach AfDipA from membrane, and even more efficiently (Fig. 5j). Consistent with this, AfDipBΔLoop showed stronger stimulation on AfDipA GTPase activity (Supplementary Figure 12f). However, without this loop, the amount of AfDipB co-sedimented with liposomes was significantly decreased (Fig. 5j). This decline of membrane binding is unlikely to result from the loss of interaction of AfDipB is unlikely to result from the loss of interaction of AfDipB with AfDipA, as under identical incubation conditions (3 µM AfDipA and 6 µM AfDipB), AfDipA only increased membrane-bound AfDipB by 20%, while the deletion of the loop caused membrane-bound AfDipB to plummet from 60% to 10% (Fig. 5j). In H. volcanii H26 cell, both mNG-DipBΔLoop and DipBΔLoop-mNG kept the pole-to-pole oscillation, while the mNG-DipBΔLoop showed shorter oscillation period, where the oscillation period of 63s increased from 9.9% for mNG-DipB to 33.5% for mNG-DipBΔLoop (Fig. 4k and Supplementary Figure 12h). However, DipBΔLoop-mNG didn’t form the ring like structure (Supplementary Figure 12h and Supplementary Movie 8). Furthermore, DipB-mNG lost the oscillation in the ΔdipB cells after deleting the loop region, suggesting that DipBΔLoop-mNG alone is not sufficient to sustain the oscillation (Supplementary Movie 8). Surprisingly, in addiction to membrane association, DipB also exhibits DNA binding activity (see Supplementary Results for more details).

**Fig. 4:**
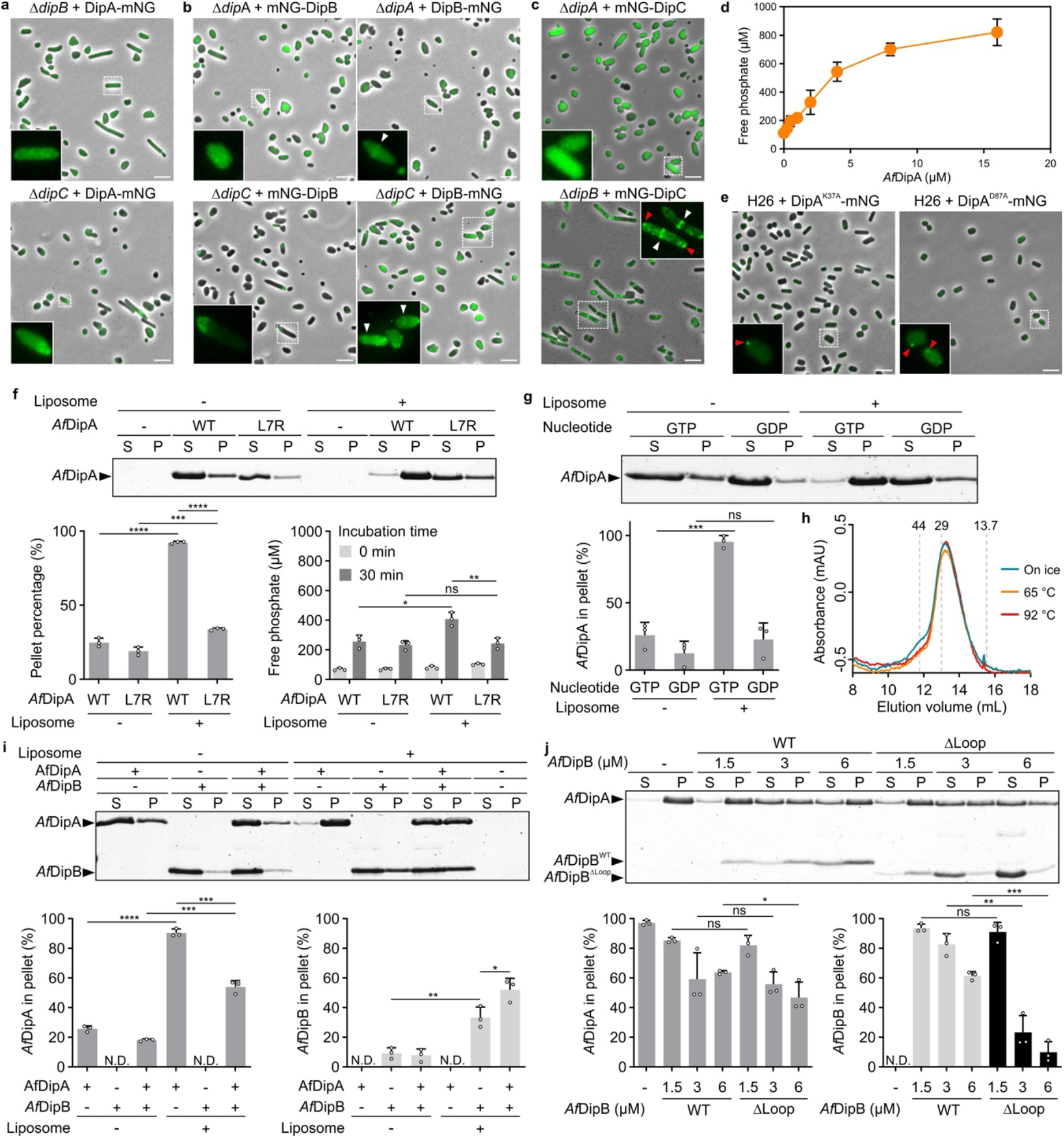
Interdependency of Dip proteins in localization and function. Fluorescence microscopy of (**a**) DipA-mNG in Δ*dipB* and Δ*dipC*, (**b**) mNG-DipB and DipB-mNG in Δ*dipA* and Δ*dipC*, (**c**) mNG-DipC in Δ*dipA* and Δ*dipB*. Representative ring-like structures are indicated by white arrowheads, foci are indicated by red arrowheads. **d**, GTPase activity of *Af*DipA. Free phosphate was quantified using the malachite green-ammonium molybdate colorimetric assay after incubating *Af*DipA at 70 °C for 40 min (mean ± SD, n=3). **e**, Fluorescence microscopy of DipA^K37A^-mNG and DipA^D87A^-mNG in *H. volcanii* H26 cells. Representative foci are indicated by red arrowheads. **f**, Liposome binding to GTPase activity of *Af*DipA. 3 µM of *Af*DipA or its MTS mutant (*Af*DipA^L7R^) were incubated with (+) or without (-) liposomes (0.5 mg/mL) at 80 °C for 30 min. Supernatant (S) and pellet (P) fractions were separated by centrifugation and analyzed by SDS-PAGE. *Af*DipA in pellet fraction and free phosphate after incubation were quantified (mean ± SD, n=3). **g**, Requirement of GTP for membrane binding of *Af*DipA. 3 µM *Af*DipA was incubated in the manner as (**f**) in the presence of 1 mM GTP or GDP, with or without liposomes (0.5 mg/mL). *Af*DipA in pellet fraction was quantified (mean ± SD, n=3). **h**, Size exclusion chromatography of *Af*DipB. 500 µL of 10 µM purified *Af*DipB (preincubated at different temperatures for 10 min) was loaded on a Superdex 75 10/300GL column. Absorption at 280 nm at the different elution volumes are shown. Elution volumes of protein size markers (in kDa) are indicated. **i**, Effect of *Af*DipB on *Af*DipA membrane binding. 3 µM *Af*DipA and 6 µM *Af*DipB were incubated with or without liposomes (0.5 mg/mL) at 80 °C for 30 min. Sedimentation assay were described above (**f)**. *Af*DipA (lower left) and *Af*DipB (lower right) in pellet fraction were quantified (mean ± SD, n=3). **j**, Impact of the *Af*DipB loop region (AA: 73-85) on *Af*DipA and *Af*DipB membrane binding. 3 µM *Af*DipA was incubated with *Af*DipB wildtype (WT) and loop region deletion mutant (ΔLoop) in the presence of liposome (0.5 mg/mL). Sedimentation assay was described above (**f**) *Af*DipA (lower left) and *Af*DipB (lower right) in pellet fraction were quantified (mean ± SD, n=3). For (**a-c**) and (**e**), cells were grown in liquid Hv-Cab medium and induced with 1 mM tryptophan for 6 h. Fluorescent channel image of cells in dashed boxes are magnified. Scale bar, 5 µm. For (**f**), (**g**), (**i**) and (**j**), significance was tested by two-tailed t-test. * P ≤ 0.05, ** P ≤ 0.01, *** P ≤ 0.001, **** P ≤ 0.0001, ns P > 0.05, n = 3.

**Fig. 5:**
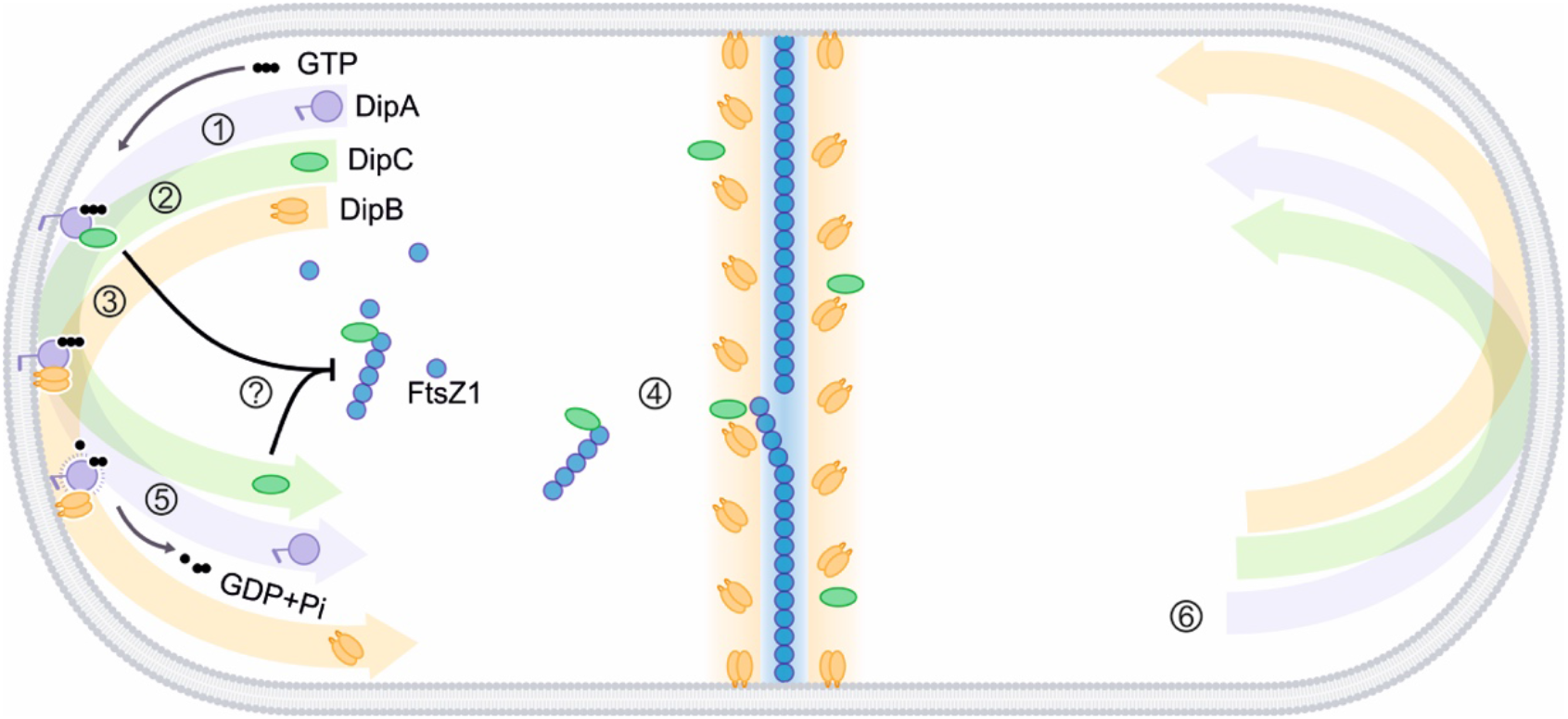
Proposed model of Dip system function in Archaea. The DipA protein first binds to the cell membrane at one of the cell poles in a GTP-bound form. DipC is then recruited by DipA to the cell pole, and enriched DipC inhibits polar FtsZ1 ring formation (by a still uncharacterized mechanism). DipA also recruits DipB to the pole, which replaces DipC. The interaction with DipB catalyzed the GTP hydrolysis by DipA, thereby converting DipA from its GTP-bound form to a GDP-bound state, which leads to the detachment of DipA from the cell membrane. Subsequently, DipA translocates to the opposite cell pole and repeats the process of sequentially recruiting DipC and DipB. Ultimately, this leads to the pole-to-pole oscillation of Dip system, resulting in polar accumulation of DipC and enabling FtsZ1 ring formation at midcell.

## Discussion

In this study, we identify a system widely conserved across Archaea that we term the Dip system. This system consists of three proteins and exhibits a strong co-occurrence with the FtsZ1 (Fig. 1b). By analyzing the phenotypes of deletion and overexpression of each gene in *H. volcanii*, combined with a detailed biochemical characterization of the individual proteins, we conclude that Dip is an oscillatory system that positions the archaeal division machinery at midcell. Based on our results, and by analogy to the bacterial Min system, we propose a model for how the Dip system controls divisome placement in Archaea (Fig. 6).

DipA binds to the cell membrane at one cell pole in its GTP-bound form, in a similar manner to ATP-bound MinD in bacteria^17^. Once localized at the cell pole, DipA recruits DipC, which, functionally similar to MinC, inhibits FtsZ1 ring formation at the pole. Following recruitment of DipC, DipB is subsequently recruited to the cell pole and displaces DipC from DipA (similarly to how MinE disrupts the MinCD interaction). The interaction with DipB enhances the GTP hydrolysis of DipA, converting it from the GTP-bound form to the GDP-bound form. As a consequence, the membrane binding affinity of DipA is substantially reduced, leading to its detachment from the membrane and translocation to the opposite pole, where the processes of membrane attachment and sequential recruitment of DipC and DipB reoccur. This results in pole-to-pole oscillation of the Dip system, creating a concentration gradient of DipC that is highest at the poles and lowest at midcell, suppressing polar FtsZ1 polymerization and directing Z-ring formation at midcell.

Several in vivo experimental findings support our proposed model. Crucially, the absence of FtsZ1 filaments in cells overexpressing DipC suggests that DipC inhibits FtsZ1 polymerization, thereby preventing formation of the FtsZ1 ring at the cell pole (Fig. 3b). Excessive intracellular level of DipC suppresses FtsZ1 polymerization throughout the cell, leading to severe defects in cell division and the formation of enlarged cells (Fig. 3c). Although overexpression of DipA in the H26 cell also disrupts FtsZ1 filament formation and causes enlarged cells, this not happened in the cells lacking DipC, indicating that DipA-mediated inhibition of cell division is dependent on DipC (Fig. 3c). However, since the purification of DipC was unsuccessful, the exact mechanism by which DipC inhibits FtsZ1 polymerization remains unclear. In E. coli, MinC interacts directly with FtsZ, and both N- and C-terminal domains inhibit FtsZ assembly, where the C-terminal domain is also required for binding to MinD^14,34,35^. DipC also contains conserved N- and C-terminal domains, which in *H. volcanii* are connected by an extremely long intrinsically disordered region (Supplementary Figure 3). Unlike MinC in *E. coli*, DipC in *H. volcanii* forms a ring-like structure at the division site that co-localizes with FtsZ1 (Supplementary Figure 9b). The coexistence of pole-to-pole oscillation and midcell ring has also been observed in cyanobacteria^36^. This observation seems inconsistent with the proposed inhibitory role of DipC on FtsZ1 ring formation. One possible explanation is that this DipC ring is highly dynamic. While DipC inhibits FtsZ1 ring formation at the cell poles, a subset of DipC remains associated with FtsZ1 and is transported to midcell as well, explaining the presence of DipC rings at the division site in *H. volcanii*. These FtsZ1-associated DipC are subsequently removed by DipB, which also localizes to the division site (Supplementary Figure 9a). This hypothesis accounts for our time-lapse time microscopy that DipC rings are much less static than FtsZ1 rings and can appear multiple times during a single division cycle (Supplementary Movie 5). This also explains why DipB forms a ring-like structure at midcell. An alternative possibility is that, similar to MinC in *B. subtilis*, DipC forms two rings flanking the FtsZ ring, thereby preventing aberrant extension of FtsZ filaments or over-initiation of FtsZ ring^37^. In B. subtilis, this midcell localization of MinC requires several other proteins, e.g. MinD, to also localizes to the division site37. However, we did not observe DipA at midcell in H. volcanii. We cannot exclude the possibility that native DipA does localize to the division site, as our localization results are derived from DipA-mNG, and the fluorescent tag may interfere with DipA localization, as suggested by the distinct localization patterns observed for mNG-DipB and DipB-mNG (Fig. 4b, c).

DipB also exhibits DNA-binding activity, in addition to its membrane-binding activity and its ability to stimulate DipA GTP hydrolysis (Supplementary Figure 12g). MinD in *E. coli* has been reported to bind DNA, leading to the hypothesis that the Min system may participate in chromosome segregation, although this idea remains controversial46. Whether the DNA-binding activity of DipB indicates a role for the Dip system in archaeal chromosome segregation remains unclear, particularly given that chromosome segregation in H. volcanii has not yet been well characterized.

Beside causing severe cell division defect, loss of the Dip system completely abolishes motility of *H. volcanii* cells (Supplementary Figure 4e, f). Motility in *H. volcanii* is regulated by MinD_4_, which also undergoes pole-to-pole oscillation^25^. It is therefore plausible that the Dip system could also influence MinD4 localization and, consequently, cellular motility. Indeed, previous studies have shown that the MinDE system in *E. coli* regulates a variety of membrane proteins involved in pattern formation, even though these proteins may be functionally unrelated^41^. Thus, in *H. volcanii*, the Dip system may simultaneously modulate the activities of FtsZ1 and MinD4, thereby coordinating cell division and motility. Given the close relationship between cell shape and motility in *H. volcanii*, the Dip system may act as an integrative regulator that spatiotemporally synchronizes cell division, shape changes, and motility.

Although the Min system and Dip system share notable functional similarities, they exhibit no homology based on sequence similarity. Why bacteria and archaea have evolved two entirely non-homologous systems to perform analogous cellular functions remains an open question. Similarly, additional studies are needed to understand whether the emergence of Min and Dip systems was driven by FtsZ-based division machinery or vice versa. Nonetheless, our discovery that Dip is an oscillatory system that positions the divisome in Archaea represents a clear example of convergent evolution for cell division placement across domains of life and is likely to stimulate future studies advancing our understanding of the origin and evolution of life.

## Material and methods

If not stated otherwise, all chemicals were either purchased from Carl Roth or Sigma.

### Growth and media

*Haloferax volcanii H26* was used as background strain and regarded as wild-type^42^, all strains used in this study are listed in Supplementary Table 1. *H. volcanii* cells were grown in either rich Hv-YPC medium (5 g/L yeast extract (Oxoid), 1 g/L Peptone (Oxoid) and 1 g/L Bacto Casamino Acids (BD Biosciences)) before plasmid transformation or selective Hv-Ca medium (5 g/L Bacto Casamino Acids (BD Biosciences))42 supplemented with an expanded trace element solution (Hv-Cab medium) after transformation^43,44^. Small liquid cultures (less than 5 mL) were grown in 15 mL tube with constant rotation at 45 °C, while large liquid cultures (e.g., 20 mL) were gown in Erlenmeyer flasks in a shaker (120 rpm) at 45 °C. Hv-YPC and Hv-Ca solid plates with 1.5% Bacto agar (BD Biosciences) were incubated in plastic boxes or bags to prevent evaporation at 45 °C. E.coli cells used for cloning were grown in lysogeny broth (LB) medium supplemented with proper antibiotics according to selection markers (100 μg/mL ampicillin or 25 μg/mL kanamycin)^45^ at 37 °C for overnight with 150 rpm shaking.

### Cloning

Plasmids used in this study were constructed with restriction enzyme-based cloning, Gibson assembly^46^, in vivo ligation^47^, and in vitro ligation methods. The enzymes used for plasmid generation were purchased from New England Biolabs (NEB) and used according to the manufacturer’s protocol. To construct the knockout plasmids, about 500 bp upstream and 500 bp downstream regions of the genes of interest were amplified from genomic *H. volcanii* DNA. The upstream, downstream fragments and linearized pTA131^42^ were then transformed into *E*.*coli* cells and ligated via in vivo ligation using 15 bp of homologous regions per site. The upstream and downstream fragments of *hvo_3012* knockout plasmid were ligated via a BamHI restriction site.

Plasmids pSVA5997 and pSVA5998 were used to construct localization plasmids for expression of C terminal or N terminal mNeonGreen tagged proteins respectively^8^. Plasmids pSVA6881 and pSVA6882 were used as the backbones to construct dual localization plasmids. The desired genes were amplified from genomic *H. volcanii* DNA and the restriction sites were integrated by correspond primers. Genes were cloned into vectors with restriction enzyme-based method.

The plasmid expressing HaloTag-DipC was constructed by changing the mNeonGreen tag of pSVA6455 into HaloTag. The plasmid expressing FtsZ1-HaloTag was generated by changing the HpaA of pTA963 HpaA-HaloTag^48^ into FtsZ1. The inserts and linearized vectors were ligated via Gibson assembly using NEB HiFi DNA assembly kit following the manufacturers’ protocol.

Plasmids for homologous protein expression in *H. volcanii* were constructed via cloning genes into backbone plasmid pTA1392^49^ with restriction enzyme-based cloning method. Heterologous His-SUMO tag protein expression plasmids were constructed with in vivo ligation by transforming gene fragments and linearized plasmid pSVA13429^8^ into *E*.*coli* cells.

Plasmids for expression of point-mutated or truncated proteins were generated by in vitro ligation method. Briefly, the backbone plasmid expressing wild-type proteins was linearized with mutation modified or truncation primers. The linearized plasmids were phosphorylated with T4 polynucleotide kinase (NEB), and then circularized by T4 DNA ligase (NEB) according to the manufacturer’s protocol.

To construct plasmids for genomic *gfp* tag integration, gene fragments containing the *gfp* tag were amplified from localization plasmids, and were ligated to downstream and linearized pTA131 by in vivo ligation.

All plasmids were verified by sanger sequencing. The plasmids, primers and restriction enzymes used in this study are listed in Supplementary Table 2 and Supplementary Table 3.

### Transformation in *Haloferax volcanii*

Plasmids were first unmethylated through a *dam*^*-*^ */dcm*^*-*^ *E*.*coli* strain, then the polyethylene glycol 600 (PEG 600) based method was used to transform *H. volcanii*^42^. The *H. volcanii* (H26 or *dip* gene knockout strains) cells were grown in 5 mL Hv-YPC medium to reach an optical density (OD_600_) of 0.8 and harvested by centrifuging at 3,000 g for 10 min. Cells were resuspended in 2 mL buffered spheroplasting solution (1 M NaCl, 27 mM KCl, 50 mM Tris-HCl (pH 8.5) and 15% w/v sucrose) and spined down at 6,000 rpm for 8 min. The cell pellet then resuspended again in another 600 μl buffered spheroplasting solution, and 200 μL suspension was used for each transformation. 50 mM EDTA (pH 8.0) was gently added to the cells and then incubated at room temperature for 10 min. 30 μl mixture of 1 μg unmethylated plasmids and unbuffered spheroplasting solution (1 M NaCl, 27 mM KCl, 83 mM EDTA (pH 8.0) and 15 % w/v sucrose) was added to the cells and was gently mixed. After 5 min incubation, 250 μL 60% PEG 600 (diluted by unbuffered spheroplasting solution) was added, gently mixed and incubated for 30 min. The spheroplasts were mixed with 1.5 mL spheroplast dilution solution (3.75 mM CaCl_2_, 23% saltwater and 15% w/v sucrose) and collected by centrifuging at 6,000 rpm for 8 min. The pellet was resuspended gently in 1 mL of regeneration solution (3 mM CaCl_2_, 18% saltwater, 1× YPC and 15% w/v sucrose) and followed by 1.5 h static and 3.5 h rotating incubation. Finally, cells were collected and resuspended in 1 mL transformant dilution solution (3 mM CaCl_2_, 18% saltwater and 15% w/v sucrose). 100 μL resuspension was plated on selective Hv-Ca plates.

### Generation of deletion and gfp.tag integrated strains

The pop-in and pop-out method was used to generate deletion strains^42^. The knockout plasmids pSVA13785, pSVA6813 and pSVA6814 were transformed into *H. volcanii* strain H26 as described above to construct *dipC, dipB* and *dipA* deletion strain respectively. One colony per transformant which expected to have integrated knockout plasmid into genome (pop-in) was transferred and incubated in 5 mL Hv-YPC medium for overnight. Subsequently, cell culture was diluted into fresh Hv-YPC medium (1:500) and grown overnight (pop-out). After three times dilution, cells were plated on Hv-Ca plates supplemented with 50 μg/mL 5-FOA and 0.09 mM uracil. Possible knockout colonies were screened via colony PCR. PCR products of positive colonies were sequenced to confirm the gene deletions. Plasmids pSVA13937 (integrate *gfp* tag to genomic *sepF*) and pSVA13958 (integrate *gfp* tag to genomic *ftsZ1*) were transformed into H26 cells followed by pop-in and pop-out steps as described above. Colonies were screened by PCR and verified via sequencing PCR products.

### Growth curves

The wild-type strain H26 and knockout strains HTQ910 (Δ*dipC*), HTQ909 (Δ*dipB*) and HTQ912 (Δ*dipA*) were transformed with empty expression plasmid pTA1392 and grew on Hv-Ca plates. For each strain, 3 individual colonies from plates were inoculated in 5 mL Hv-Cab medium and grew overnight. The pre-cultures were then diluted in 20 mL fresh Hv-Cab medium to an OD_600_ of 0.02. The growth curves of 20 mL cultures were automatically measured with the cell growth quantifier (CGQ, Software (CGQuant 7.4)) from Aquila Biolabs GmbH at 45 °C and 120 rpm shaking.

### Spot survival assays

The spot dilution assay was performed to test the viability of deletion strains. Colonies with empty expression plasmid pTA1392 were picked and grew in 5 mL Hv-Cab medium for overnight. The pre-cultures were then diluted with fresh Hv-Cab medium to an OD_600_ of 0.2. Those cultures were serially diluted 10^−6^ times, and 5 μl of each dilution per strain was dropped on Hv-Ca plates. Those plates were incubated for 2 days at 45 °C in sealed plastic bags. For each strain, 3 biological replicates were performed.

### Motility assays

The semi-solid plates were prepared by dissolving 0.33% agar in Hv-Ca medium. The pre-cultures with an OD_600_ about 0.3 were prepare with Hv-Cab medium and stabbed on the semi-solid plates. After 4 days incubation at 45 °C, the plates were scanned and the diameter of the motility ring was measured. For each strain, 3 technical and 3 biological replicates were performed. Motility assays to check for the functionality of mNeonGreen tagged proteins or non-tagged proteins expressed from plasmids were performed as described above except adding 1 mM tryptophan. The diameter of the motility ring was measured by ImageJ (v5.14f)^50^.

### Microscopy and image analysis

Phase contrast and fluorescence light microscopy was used to investigate the cell shape and protein localization in *H. volcanii* cells. Colonies with expression plasmids were picked and inoculated in 5 mL Hv-Cab medium. After overnight incubation, the pre-cultures were diluted into 20 mL fresh Hv-Cab medium and grown to an OD_600_ about 0.02. For the cells contained plasmids with tryptophan inducible promoter, 0.25 mM or 1 mM tryptophan was added to the medium to induce expression. Before imaging, 5 µL cell culture was spotted on a 1% w/v agarose pad containing 18% saltwater (144 g/L NaCl, 18 g/L MgCl_2_·6H_2_O, 21 g/L MgSO_4_·7H_2_O, 4.2 g/L KCl and 12 mM Tris-HCl, pH 7.5) on a glass slide. Sample was dried in room temperature and covered with a glass coverslip. Images were acquired with an inverted microscope at 100 x magnification (Zeiss Axio Observer.Z1, controlled via Zeiss Blue v.3.3.89).

To acquire oscillation movies, the microscope chamber was heated at 45 °C and images were captured every 15 s for 5 min or 10 min. The overnight time-lapse imaging was performed using the CellASIC ONIX2 microfluidic system and B04A-03 plates. The microfluidic channel was first washed with ddH_2_O for 3 min, PBS buffer (580 mM Na_2_HPO_4_, 170 mM NaH_2_PO_4_ and 680 mM NaCl, pH 7.3) for 3 min and fresh medium for 5 min. For cell loading, cell cultures with an OD_600_ about 0.1 were flowed for 15 s into the chamber at 13.8 kPa. Cell images were acquired every 15 min for 16 h at 45 °C with a constant flow of fresh medium at 5 kPa.

To determine the cell area (length), microscopy images were analyzed with FIJI (v1.54f)^50,51^ program and the MicrobeJ (v.5.13l)^52^ plug-in. Briefly, cell outlines were detected according to phase-contrast image by default segmentation method, with the medial axis mode, area > 0.3 µm^2^ and exclude-on-edges options, then manually corrected. To analyze localization of DipC and SepF, the cell outlines were detected as described above according to phase-contrast channel image, the fluorescence maxima were detected using the Foci and Basic modes with an area of > 0.02 according to fluorescent channel image. The demography was plotted in ResultJ by normalizing the intensity and sorting along the cell length. Cell morphology and fluorescent maxima data were exported and visualized in GraphPad Prism (v10.2.3).

The oscillation time stack images were first aligned using MultiStackRegistration (v1.46.2)^53^ in rigid body mode according to the phase-contrast image. The images were then transformed into 8-bit pictures and the cells were segmented from the first frame of the phase-contrast channel image using automated thresholding (Otsu) and watershed segmentation, followed by particle analysis. The segmented cells were manually curated to ensure accurate single-cell ROIs. For each accepted cell, geometric parameters (length, width, centroid, and orientation) were measured, and cells below a minimum length threshold (10 pixels) were excluded. Two circular pole regions (diameter = 8 pixels) were defined along the major axis based on cell length and centroid position. Mean fluorescent intensity at both poles was measured separately across all frames, and the pole1-to-pole2 intensity ratio was calculated over time. Cell morphological data (length, width, centroid, angle) and temporal pole intensity ratio values were exported. To calculate the oscillation period, pole intensity ratio values were linearly detrended using pracma package (v2.4.4)^54^ in R (v4.3.3)^55^. A Fast Discrete Fourier Transform by ftt() function from stats package (v4.4.3)^55^ was then applied to the detrended data to identify the dominant oscillation frequency, from which the primary period was calculated. The oscillation period was export for further visualization in GraphPad Prism (v10.2.3).

### 3D-structural illumination microscopy (3D-SIM) and single particle tracking (SPT)

Slides and coverslips used for imaging were cleaned with 1 M KOH overnight, rinsed with deionized H_2_O and dried with pressurized air. Agar pads where prepared with the aid of gene frames (Thermo Fisher) and 1% (w/v) low melting agarose (agarose, low gelling temperature, Sigma-Aldrich) in sterile filtered Hv-Cab media (0.2 μm pores).

*H. volcanii* H26 and mutant strains were grown in Hv-Cab to an OD_600_ of approximately 0.1. The strains expressed proteins of interest with a HaloTag fusion ectopically with a tryptophan inducible promoter; 0.25 mM or 1 mM tryptophan was added to the medium to induce expression. 1 ml of culture was stained with 30 nM HaloTag R110-Direct Ligand (Promega) for 30 min at 45 °C at 600 rpm on a standard 2 ml heat block (Eppendorf). Cells were harvested (6000 rpm, 2 min, RT) and washed three times in filtered Hv-Cab media. Cells were then spotted on cleaned slides prepared with agarose pads, allowed to dry at room temperature and covered with a coverslip.

3D-SIM and SPT were performed with an Elyra 7 (Zeiss) with lattice SIM^2^ inverted microscope equipped with pco.edge sCMOS 4.2 CL HS cameras (PCO AG), connected through a DuoLink (Zeiss). Cells were observed through an alpha Plan-Apochromat 63×/1.46 Oil Korr M27 Var2 objective in combination with an Optovar 1 ×, and 1.6 × (Zeiss) magnification changer, yielding a pixel size of 97 nm and 63 nm for SPT and 3D-SIM respectively. During image acquisition, the focus was maintained with the help of a Definite Focus.2 system (Zeiss). Fluorescence was excited with a 488 nm (100 mW, 50%) laser, and signals were observed through a multiple beam splitter (405/488/561/642 nm) and laser block filters (405/488/561/642 nm) followed by a Duolink SR DUO (Zeiss) filter module (emission filter BP 495-590).

3D-SIM images were acquired in a Z stack consisting of 13 slices 101 µm apart, and 15 phases were used. Images were reconstructed using the 3D SIM^2^ module integrated in the ZEN 3.0 SR (black edition) software (Zeiss). Final 3D-SIM images were obtained by generating maximum projections of all the Z-slices obtained. FIJI^51^ was used to convert image stacks into maximum projections and 3D projections. 3D projections were made using the 3DSuite plugin (v4.1.7)^56^.

For each SPT time lapse series, 10000 frames were taken with 20 ms exposure time (∼24 ms with transfer time included) and 50% 488 nm intensity laser. Images were acquired in TIRF mode (63° angle). The online SMLM module in the ZEN 3.0 SR (black edition) was used to identify individual spots per frame with a signal to noise ratio of 6 including overlapping spots, and exported as tables. The tracks were reconstructed using TARDIS^57^ followed by swift tracking software (version 0.4.3) (Endesfelder et al, manuscript in prep; Swift version 0.4.3 and all subsequent versions of the swift software can be obtained on the swift beta-testing repository (http://bit.ly/swifttracking)). Swift parameters were obtained from TARDIS using the tabular data in thunderSTORM format with default settings (maximum jump distance of 3 μm, Δt of 3 frames) with slow and exact background subtraction accuracy. The populations were auto-chosen. The swift GUI was run using the obtained parameters to connect individual spots into tracks. The data was further visualized in R (v4.5.0)^55^ with ggplot2 (4.0.1)^58^.

### DNA staining in *H. volcanii*

Single colonies of H26, ΔdipA, ΔdipB and ΔdipC strains were picked and inoculated in 5 mL Hv-Cab medium as pre-cultures. 50 µg/mL uracil was added to complement the uracil auxotrophy. After overnight incubation, the pre-cultures were diluted into 20 mL fresh medium and grown to an OD_600_ about 0.1. 1 mL cell culture for each sample were taken into 1.5 mL tubes and incubated with 5 µM nucleic acid dye SYTO 13 Green (Thermo Fisher Scientific) for 10 min. After incubation, 5 µL cell culture was spotted on a 1% w/v agarose pad and imaged with microscope as described above.

### Hetereologous expression and purification of A. fulgidus SepF, DipA and DipB

*Af*SepF protein was expressed and purified as previously reported with minor modifications^8^. Briefly, *Af*SepF was overproduced in *E. coli* strain Rosetta (DE3) by inducing expression from plasmid pSVA13587 with 0.5 mM isopropyl-β-D-1-thiogalactopyranoside (IPTG) in 2 liters LB medium supplemented with 100 µg/mL kanamycin at 37 °C for 3 h. Cells were harvested and resuspended in buffer A (50 mM Na_2_HPO_4_, 300 mM NaCl, adjusted with NaH_2_PO_4_ to pH 8.0) containing 10 mM imidazole and lysed by three passes through a French press at 1,000 psi. Cell debris was removed by centrifugation at 6,800 g for 10 min (JA25.50 rotor; Beckman Coulter). To remove *E. coli* proteins, the cell lysate was incubated at 70 °C for 20 min under constant shaking, followed by centrifugation at 21,000 g for 10 min at 8 °C. A final centrifugation step was performed at 83,540 g for 45 min at 4 °C before loading the supernatant on a 5 mL HisTrap HP column (Cytiva), which was pre-equilibrated with buffer A. The column was connected to an Äkta purifier (Cytiva) operated with the Unicorn software (v.5.11). After washing with 5 column volumes (CV) of buffer B (300 mM NaCl, 20 mM imidazole and 50 mM Na_2_HPO_4_, pH 8.0), proteins were eluted with buffer C (300 mM NaCl, 250 mM imidazole and 50 mM Na_2_HPO_4_, adjusted with NaH_2_PO_4_ to pH 8.0). The 6 mL peak fractions were incubated overnight at 4 °C with 2.5 µg/mL SUMO protease in the presence of 2 mM dithiothreitol (DTT) and 0.1% NP-40. The digested samples were further purified by size-exclusion chromatography using a HiLoad 16/600 Superdex 75 pg column (Cytiva) equilibrated with buffer D (150 mM NaCl and 25 mM Tris-HCl, pH 8.0).

*Af*DipA and its L7R mutant were expressed in *E. coli* strain Rosetta (DE3) with plasmids pSVA7403 and pSVA7459, respectively. Cells were grown in 2 liters LB medium supplemented with 100 µg/ml kanamycin and 0.5 mM IPTG at 37 °C for 3 h. Cells were harvested and resuspended in buffer A containing 10 mM imidazole and lysed by three passes through a French press at 1,000 psi. Cell debris was removed by centrifugation at 6,800 g for 10 min (JA25.50 rotor; Beckman Coulter). To remove *E. coli* proteins, the cell lysate was incubated at 70 °C for 20 min under constant shaking, followed by centrifugation at 8 °C for 10 min at 21,000 g. A final centrifugation step was performed at 4 °C for 45 min at 83,540 g before loading the supernatant on a 5 mL HisTrap HP column (Cytiva), which was pre-equilibrated with buffer A. The column was connected to an Äkta purifier (Cytiva) operated with the Unicorn software (v.5.11). After washing with 5 CV of buffer B, proteins were eluted with buffer C. The 6 mL peak fractions were incubated at 4 °C for 3 hours with 2.5 µg/mL His-tagged SUMO protease in the presence of 2 mM dithiothreitol (DTT) and 0.1% NP-40. The buffer was then exchanged to SEC buffer (100 mM HEPES-KOH pH 7.4, 150 mM KCl, and 10% Glycerol) using HiPrep 26/10 Desalting column (Cytiva), followed by Ni-affinity purification to remove His-SUMO tag and His-tagged SUMO protease.

*Af*DipB and its Δloop mutant were overproduced in *E. coli* strain Rosetta (DE3) by inducting expression from plasmid pSVA7402 and pSVA7458 using 0.5 mM IPTG in 2 liters fresh LB medium supplemented with 100 µg/mL kanamycin at 37 °C for 3 h. Cells were harvested and resuspended in Ni-NTA buffer A (100 mM HEPES-KOH and 150 mM KCl, pH 7.4) containing 10 mM imidazole and lysed by three passes through a French press at 1,000 psi. Cell debris was removed by centrifugation at 6,800 g for 10 min (JA25.50 rotor; Beckman Coulter). To remove *E. coli* proteins, the cell lysate was incubated at 70 °C for 20 min under constant shaking, followed by centrifugation at 8 °C for 10 min at 21,000 g. A final centrifugation step was performed at 83,540 g for 45 min at 4 °C before loading the supernatant on a 5 mL HisTrap HP column (Cytiva) which was previously equilibrated with buffer A. The column was connected to an Äkta purifier (Cytiva) operated with the Unicorn software (v.5.11). After washing with 5 CV of Ni-NTA buffer A supplemented with 20 mM imidazole, proteins were eluted with Ni-NTA buffer B (100 mM HEPES-KOH, 150 mM KCl, and 400 mM imidazole, pH 7.4). Proteins were purified by a HiLoad 16/600 Superdex 75 pg column (Cytiva) equilibrated with buffer E (150 mM KCl and 20 mM Tris–HCl, pH 8.0). The peak fractions were incubated for 3 hours at 4 °C with 2.5 µg/mL His-tagged SUMO protease in the presence of 2 mM dithiothreitol (DTT) and 0.1% NP-40, followed by Ni-affinity purification to remove His-SUMO tag and His-tagged SUMO protease. Samples were further purified by Superdex 75 10/300 GL column (Cytiva).

Purified proteins were frozen in liquid nitrogen and stored at -80 °C until use.

### Malachite green assay

To assess the NTP hydrolysis activity of *Af*DipA, the purified protein was incubated in a 25 µL reaction mixture containing 100 mM HEPES-KOH (pH 7.4), 150 mM KCl, 10% glycerol, 5 mM MgCl_2_ and 1 mM NTP (ATP, CTP, or GTP) at 70 °C for 40 min. After the indicated incubation time, 2 µL incubated reaction mixture was diluted into 20 µL Milli-Q water, then immediately quenched on ice. The diluted mixture was mixed with 180 µL malachite green-ammonium molybdate solution in a 96-well plate and incubated at room temperature for 20 min. Absorbance at 620 nm was measured using CLARIOstar plate reader (BMG LABTECH). Free phosphate concentration was calculated by a standard curve generated with KH_2_PO_4_.

### *E*.*coli* liposome preparation

Commercial *E. coli* polar lipid extract (Avanti Polar Lipids) was further purified by acetone/ether washing as previously described^59^, the purified lipids were stocked in chloroform. Before using, the lipids were dried slowly in a rotary evaporator, washed with ethanol, dried again and resuspended in liposome buffer (20 mM HEPES-KOH pH 7.4, 100 mM KCl, and 2 mM DTT) to a concentration about 20 mg/mL. The lipid suspension was sonicated for 15 min in a water bath, frozen in liquid nitrogen and allowed to thaw slowly at room temperature. This freeze-thaw cycle was repeated four times. Aliquots of the liposome were frozen by liquid nitrogen and stocked in -80 °C.

### Liposome sedimentation assay

The sedimentation assay was performed as previously described with minor modifications^17^. Briefly, *Af*DipA and/or *Af*DipB, or their mutants were incubated in a reaction mixture containing 50 mM HEPES-KOH (pH 7.4), 100 mM KCl, 5 mM MgCl_2_, and 1 mM GTP in the presence of 0.5 mg/mL *E. coli* liposomes at 80 °C for 30 min or indicated incubation time. Liposomes were sedimented by centrifugation at 20,000 g for 1 min at room temperature. The supernatant was collected separately, and the pellet was resuspended in an equal amount of reaction mixture lacking GTP. Both fractions were analyzed by 15% SDS-PAGE followed by Coomassie Brilliant Blue staining. To quantify the percentage of the precipitated fraction, densitometric analysis was performed on defined regions of interest (ROIs). Background-subtracted peak areas corresponding to the protein bands were measured, and the proportion of the precipitated fraction was calculated by dividing its intensity by the sum of the intensities of the precipitated and supernatant fractions. To monitor free phosphate concentration changes, 2 µL aliquots were taken and analyzed using the Malachite Green assay as described above.

### Electrophoretic mobility shift assay

277-bp DNA fragment was amplified by PCR using plasmid pTA131 as template and primers pTA131_fw_seq and pTA131_rev_seq^60^. The indicated amount of *Af*DipB and *Af*SepF were incubated with 30 nM 277-bp DNA at 37 °C (or indicated temperature) for 10 min in 10 μL reaction mixture containing 20 mM HEPES-KOH (pH 7.4), 50 mM KCl, 0.1 mg/mL BSA, and 7% glycerol. The samples were then analyzed by 5% native PAGE at 100 V for 40 min in Tris-Borate buffer and stained with HDGreen Plus DNA stain (INTAS). Gel images were acquired using the FAS-Digi (NIPPON Genetics).

### Bioinformatic analysis

766 archaeal genomes were retrieved from NCBI genome database as of May 2025 (Supplementary Data 1) which represented the major archaeal phyla. The archaeal phylogenetic tree was built by the program GToTree (v1.8.14)^61^, using the prepackaged 76 archaeal single-copy genes (-H Archaea). Briefly, the protein products files (.faa file) were used as input. Target proteins were identified with HMMER3 (v3.3.2)^62^, individually aligned with Muscle (v5.1)^63^, trimmed with TrimAl (v1.4.rev15)^64^, and concatenated prior to phylogenetic estimation with FastTree2 (v2.1.11)^65^. The Pfam domain PF18822^66^ corresponding to the CdvA was also searched with GToTree (- p).

The MacSyFinder (v2.1.3)^32^ was used to investigate the distribution of dip systems in Archaea. A MacSyFinder model (Supplementary Data 4) was built with the HMM profiles (PF09845 for *dipC*, PF09846 for *dipB*, and PF01926 for *dipA*) retrieved from the Pfam database as of May 2025^66^. All three genes were set as mandatory gene with at least 2 mandatory genes required, and the maximum inter gene space was set as 1. Before the scanning, the protein products files were reordered according to the genomic feature files (.gff file). Then all 766 archaeal genomes were scanned by MacSyFinder2 in ordered replicon mode (--db-type ordered_replicon) with default setting except a profile coverage of 30% (--coverage-profile 0.3). The results were manually checked and the hits identified as best solution by MacSyFinder for each genome were extracted and analyzed. The gene coordinates were retrieved from the genomic feature files and merged to the MacSyFinder results (Supplementary Data 2) The multiple sequence alignment of Dip proteins was achieved by Clustal Omega (v1.2.4)^67^ and visualized by ENDscript (v3.2)^68^.

To find the FtsZ1 and FtsZ2 homologs, 46 FtsZ1 and 36 FtsZ2 aligned protein sequences^5^ were collected and used to build custom HMM models with hmmbuild in HMMER package (v3.4)^62^. The hmmsearch from the HMMER package (v3.4)^62^ was used to search the FtsZ1 and FtsZ2 homologs in all archaeal genomes with default parameters except for an E-value threshold ≤ 1e-10. The identified homologs were manually checked to discard false positives according to published information^4,5,7^. Results were mapped onto the phylogenetic tree using iTOL (v7)^69^ and R (v4.4.3)^55^ package ggtree (v3.10.1)^70^ and gggenes (v0.5.1)^71^.

Protein 3D structures were predicted by Alphafold 3^30^ and visualized with UCSF ChimeraX (v1.9)^72^. The proteins structures were aligned by UCSF ChimeraX matchmaker tool with Needleman-Wunsch alignment algorithm and default parameters. Liquid-liquid phase separation potential of protein was predicted by FuzDrop tool^31^. Helical wheel diagram for the predicted amphipathic helix was done using Heliquest^73^.

### Statistics and graphics

For overnight microscopy, at least two independent videos from distinct replicates were recorded, showing essentially the same results. All other experiments were at least performed in three independent biological replicates, and all replication attempts were successful. If not stated otherwise, the data measurements were analyzed and visualized with GraphPad Prism (v10.2.3). In general, for comparisons of two groups, significance was determined by two-tailed, unpaired Student’s t-test. P values < 0.05 were considered statistically significant. The model and final figures were assembled with CorelDRAW 2018.

## Data availability

The data that support the findings of this study are available in the Supplementary Information, Supplementary Data and Source Data. Source data are provided with this paper.

## Authors contributions

W.H. and S.V.A. conceived and designed the project. W.H performed the bioinformatics analysis, constructed most of the plasmids and strains, and performed the primary in vivo experiments. R.Y constructed the heterologous expression plasmids, purified the proteins of *A. fulgidus*, and performed all in vitro experiments. P.N: constructed plasmids for DipC knockout and localization, and strains with genomic integrated *gfp* tag. K.P. constructed the plasmids, and performed experiments for the 3D-SIM and SPT. S.S contributed to the phylogeny analysis. C.v.d.D. contributed to the protein purification and liposome sedimentation analysis. W.H. recruited the data, assembled the figures and wrote the manuscript. M.B and S.V.A reviewed drafts of the article. S.V.A supervised the work and acquired funding.

## Acknowledgements

We thank Michal Rössler for building and drawing the final model. We thank Martin Loose at Institute of Science and Technology Austria (ISTA) for valuable suggestions on liposome sedimentation analysis. We thank Iain G. Duggin at University of Technology Sydney for providing the dual expression plasmid of FtsZ2-GFP and FtsZ1-mCherry (pIDJL134). We thank Xing Ye (ISTA) for providing the His-SUMO expression plasmid pSVA13429. We would like to thank the developers of the MultiStackRegistration project (https://github.com/miura/MultiStackRegistration) for providing valuable open-source tools. Molecular graphics and analyses performed with UCSF ChimeraX, developed by the Resource for Biocomputing, Visualization, and Informatics at the University of California, San Francisco, with support from National Institutes of Health R01-GM129325 and the Office of Cyber Infrastructure and Computational Biology, National Institute of Allergy and Infectious Diseases. Wei He was supported by the CSC fellowship (China Scholarship Council), Ryusei Yoshida was found by Daiichi Sankyo Foundation of Life Science. ShS received funding from the SFB 1381 (German Research Foundation, DFG) under project no. 403222702-SFB 1381). PN received funding from the VW Foundation from a Momentum grant (94993) to SVA. Funded by the European Union (ERC, ARCHCELLORG, 101142324). Views and opinions expressed are, however, those of the author(s) only and do not necessarily reflect those of the European Union or the European Research Council. Neither the European Union nor the granting authority can be held responsible for them. SVA received funding from the DFG under Germany’s Excellence Strategy (CIBSS-Exc_2189-Project ID 390939984). INST Elyra – We thank Dr. Giacomo Giacomelli (Kiel University) for help with the single-molecule localization microscopy.

## Notes

### Competing Interest Statement

The authors have declared no competing interest.

