## Supplementary Information for "A conserved oscillatory system that positions the divisome in Archaea"

[*] These authors contributed equally

### Supplementary Results and Discussion

#### Functionality of mNeongreen-fusion proteins

Some of the mNeongreen-fusion proteins were unstable or did not fully complement the deletion mutants. Below we discuss these tests in detail:

Unlike DipA-mNG, mNG-DipA was highly unstable, which resulted in low signal to noise ratio and unrecognizable oscillation. Nevertheless, small foci could still be identified (**Supplementary Movie 4**). Expression of either mNG-DipA or DipA-mNG did not lead to the formation of minicells or enlarged cells, unlike the expression of untagged DipA in H26 (**Fig. 3a**), suggesting that the mNG-fused DipA proteins are not fully functional. Consistent with this, overexpression of mNG-DipA in H26 did not affect its motility while DipA-mNG overexpression reduced the motility slightly. The expression of mNG-DipA or DipA-mNG in Δ*dipA* could also not restore the motility defect **Supplementary Figure 9c**). DipA-mNG did not oscillate when expressed in Δ*dipA* cells, suggesting that DipA functions as an oligomer (**Supplementary Movie 9**). The fusion of mNG at C-terminal inhibited the interaction of DipA with other proteins but did not supress the interaction of DipA-mNG with native DipA, so that DipA-mNG could oscillate in H26.

In agreement, the overexpression of DipB-mNG in H26 cells disturbed cell division and resulted in formation of minicells (**Supplementary Movie 10**). Motility assays also support that DipB-mNG was more functional than mNG-DipB (**Supplementary Figure 9d**). In addition, mNG-DipB did not oscillate when expressed in Δ*dipB* cells, while the ring formation and oscillation of DipB-mNG were unaffected in these cells (**Supplementary Movie 9**).

DipC-mNG also formed foci in cells, but those foci were much more static, suggesting that they were aggregates due to overexpression (**Supplementary Movie 4**). Neither mNG-DipC nor DipC-mNG overexpression in H26 produce minicells, suggesting that they were not fully functional (**Supplementary Movie 4**). mNG-DipC could still oscillate in Δ*dipC* cells, suggesting that its oscillation was not dependent on native DipC protein (**Supplementary Movie 9**). DipC-mNG slightly reduced the motility of H26 and significantly recovered the motility defect of Δ*dipC* cells (**Supplementary Figure 9e, f**). However, considering the strongly diffused signal of DipC-mNG (**Supplementary Movie 4**), those effects might be achieved by the tag cut DipC-mNG.

#### DNA binding of DipB

Electrophoretic mobility shift assays (EMSAs) confirmed that *Af*DipB can bind double-stranded DNA (dsDNA) (**Supplementary Figure 12g**). We also purified *A. fulgidus* SepF (*Af*SepF) as described previously[1], and checked whether it can bind DNA with EMSAs (**Supplementary Figure 12a**). Although the high structure similarity of

DipB and SepF, *Af*SepF didn't show DNA binding property (**Supplementary Figure 12g**). To rule out the possibility that this DNA binding of *Af*DipB was caused by *E. coli* protein contamination, we incubated the reaction mixture at high temperature (92 ℃) for 10 minutes to inactivate the *E. coli* proteins. The results showed that even after high-temperature incubation, *Af*DipB still exhibited DNA binding activity (**Supplementary Figure 12g**). The electrophoretic mobility shift assay of the *Af*DipB$^{\Delta Loop}$ indicates that the loop crucial for *Af*DipB membrane association is also essential for *Af*DipB DNA binding (**Supplementary Figure 12g**).

### Supplementary Figures

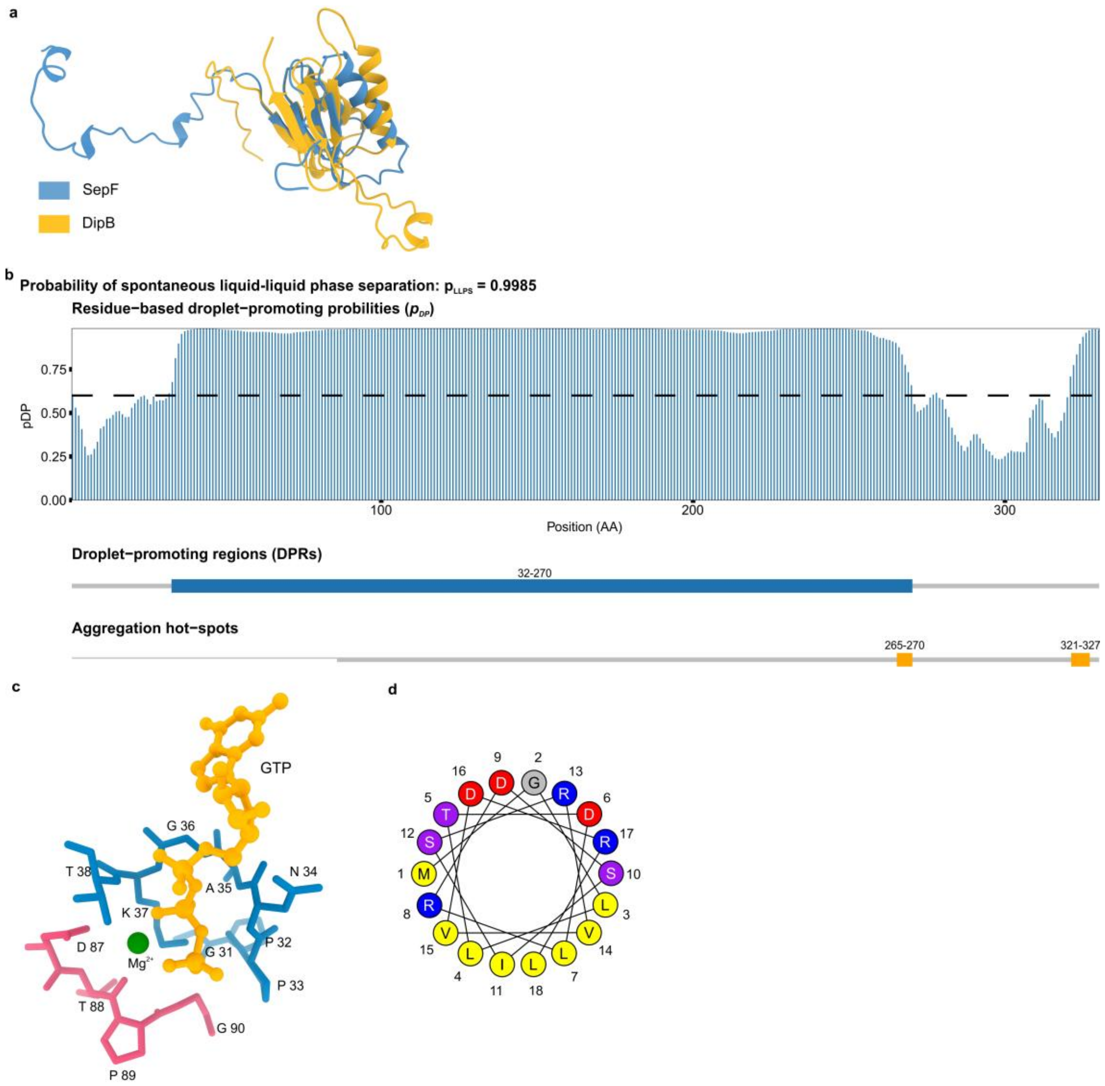

**Supplementary Figure 1: Structure features of Dip proteins.**

**a**, Structural alignment of DipB with SepF. Proteins structures were predicted with AlphaFold 3[2] (blue: SepF, orange: DipB). Structures were aligned by UCSF ChimeraX[3] matchmaker tool with Needleman-Wunsch alignment algorithm and default parameters. The root mean square deviation (RMSD) between 25 pruned atom pairs is 0.971 Å (across all 87 pairs: 14.430 Å). **b**, Evaluation of the probability of DipC to undergo liquid-liquid phase separation by FuzDrop[4] method. Top panel represents the sequence distribution of residue-based droplet-promoting probabilities ($p_{DP}$), residues with $p_{DP} \geq$ 0.60 (indicated as black dash line) is capable to promote liquid-liquid phase separation. The droplet-promoting region (DPR) ($p_{DP} \geq 0.60$) is shown by blue bar. Aggregation hot-spots that drive aggregation of condensates are displayed by orange boxes. **c**, Walker A (blue, residues 31-38, GPPNAGKT) and Walker B (red, residues 87-90, DTPG) motif atomic structure of DipA, structures were predicted with AlphaFold 3[2].

GTP is indicated as yellow ball model, $Mg^{2+}$ is represented by a green ball, amino acids are labeled with single letter code. **d**, Helical wheel diagram for the predicted amphipathic helix at the N-terminus of DipA using Heliquest[5].

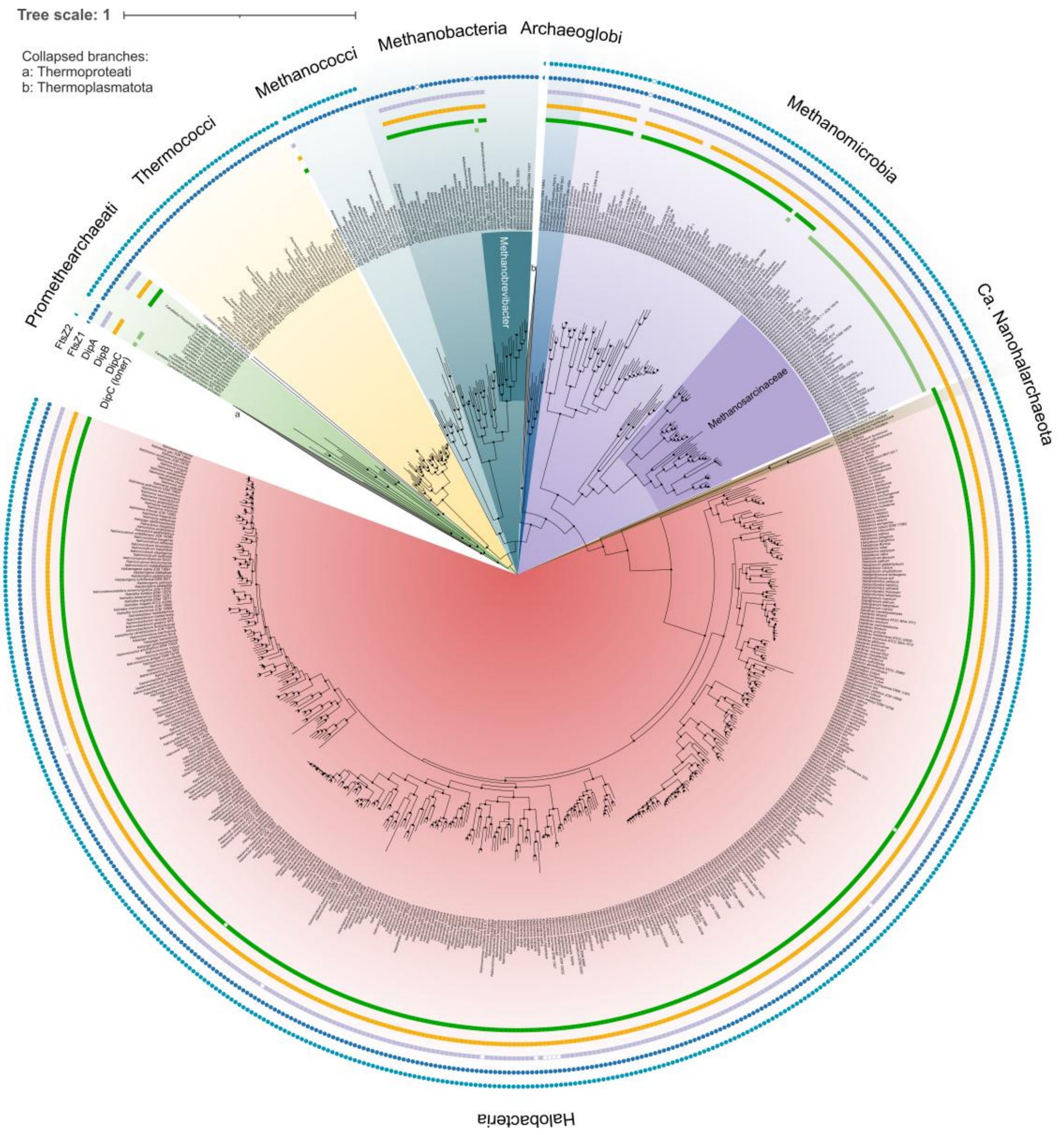

**Supplementary Figure 2: Phylogenetic distribution of Dip and FtsZ proteins in Archaea.**

Distribution of loner DipC (light green square), DipC (green square), DipB (orange square), DipA (purple square), FtsZ1 (dark blue circle) and FtsZ2 (light blue circle) were mapped to a phylogenetic tree of Archaea (Thermoproteati and Thermoplasmatota were collapsed, re-rooted at midpoint), which was generated with 766 archaeal genomes. Proteins from pseudo genes are indicated as non-filled squares or circles, semicircles indicated that the corresponding protein couldn't be identified in all the taxa of this clade. The local support values (LSV) based on the Shimodaira-Hasegawa (SH) test larger than 0.9 are indicated by dark dots. Tree scale as indicated.

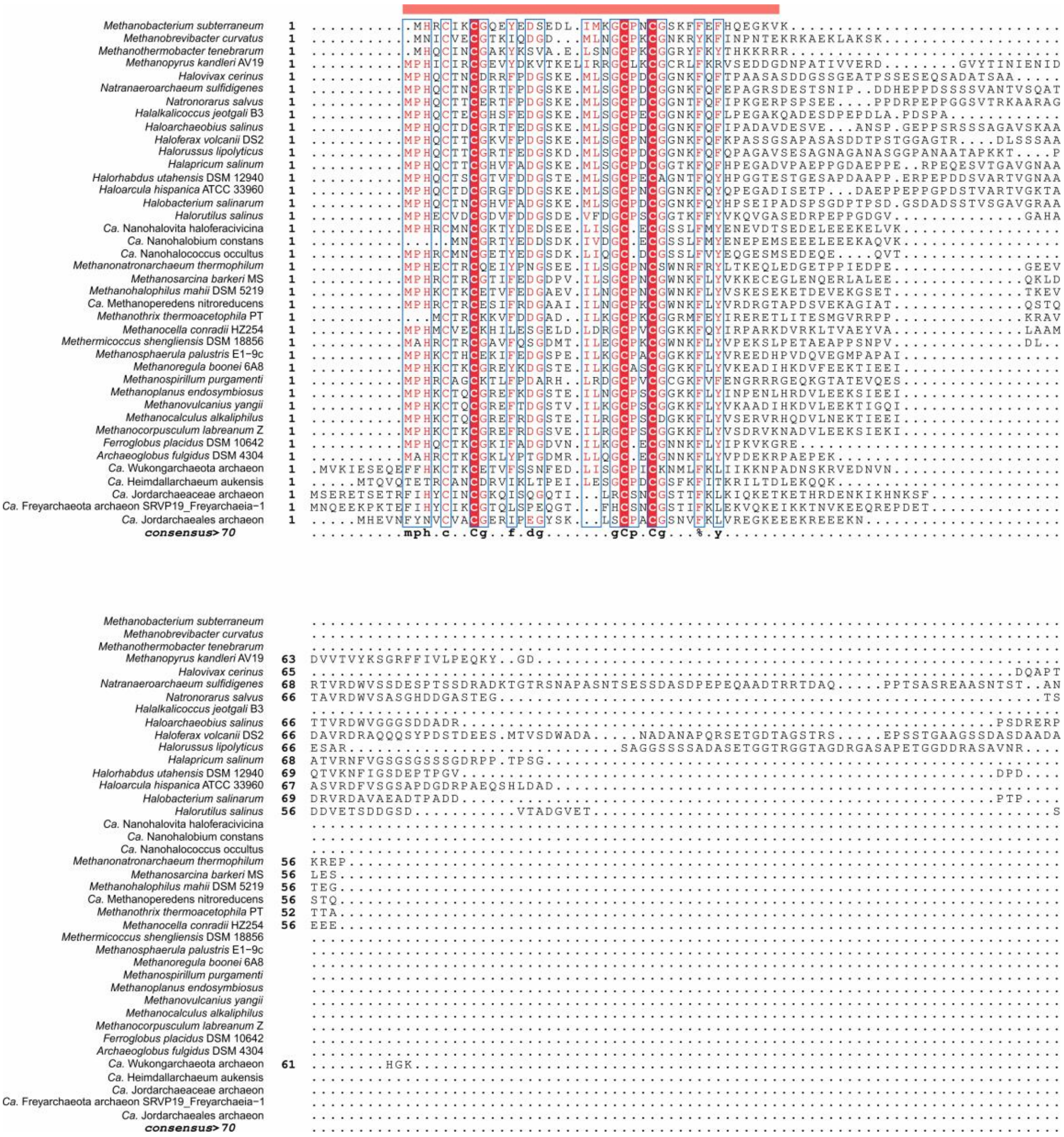
Methanobacterium subterraneum
Methanobrevibacter curvatus
Methanothermobacter tenebrarum
Methanopyrus kandleri AV19
Halovivax cerinus
Natranaeroarchaeum sulfidigenes
Natronorarus salvus
Halalkalicoccus jeotgali B3
Haloarchaeobius salinus
Haloferax volcanii DS2
Halorussus lipolyticus
Halapricum salinum
Halorhabdus utahensis DSM 12940
Haloarcula hispanica ATCC 33960
Halobacterium salinarum
Halorutilus salinus
Ca. Nanohalovita haloferacivicina
Ca. Nanohalobium constans
Ca. Nanohalococcus occultus
Methanonatronarchaeum thermophilum
Methanosarcina barkeri MS
Methanohalophilus mahii DSM 5219
Ca. Methanoperedens nitroreducens
Methanothrix thermoacetophila PT
Methanocella conradii HZ254
Methermicoccus shengliensis DSM 18856
Methanosphaerula palustris E1-9c
Methanoregula boonei 6A8
Methanospirillum purgamenti
Methanoplanus endosymbiosus
Methanovulcanius yangii
Methanocalculus alkaliphilus
Methanocorpusculum labreanum Z
Ferroglobus placidus DSM 10642
Archaeoglobus fulgidus DSM 4304
Ca. Wukongarchaeota archaeon
Ca. Heimdallarchaeum aukensis
Ca. Jordarchaeaceae archaeon
Ca. Freyarchaeota archaeon SRVP19_Freyarchaeia-1
Ca. Jordarchaeales archaeon
consensus>70

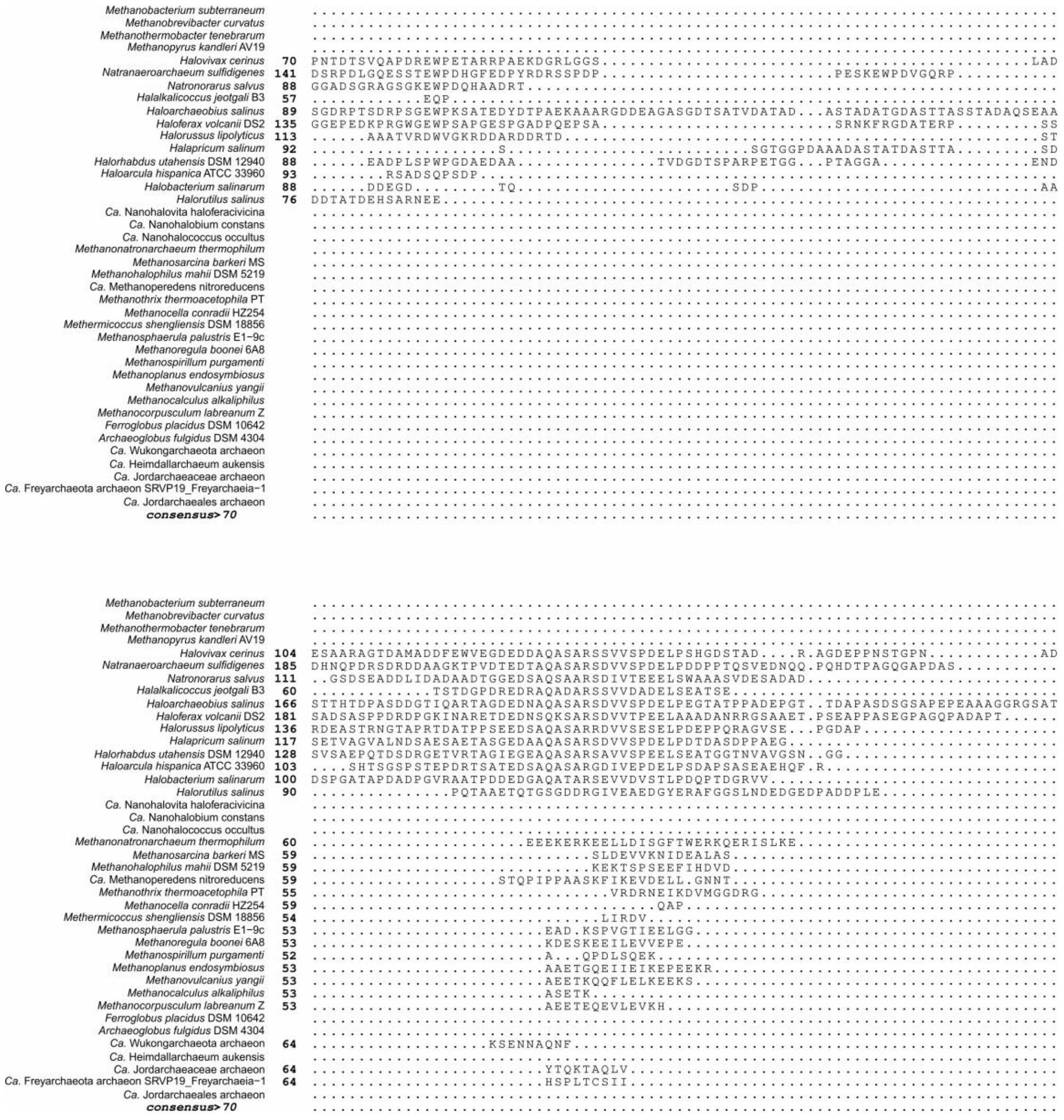
Methanobacterium subterraneum
Methanobrevibacter curvatus
Methanothermobacter tenebrarum
Methanopyrus kandleri AV19
Halovivax cerinus
Natranaeroarchaeum sulfidigenes
Natronorarus salvus
Halalkalicoccus jeotgali B3
Haloarchaeobius salinus
Haloferax volcanii DS2
Halorussus lipolyticus
Halapricum salinum
Halorhabdus utahensis DSM 12940
Haloarcula hispanica ATCC 33960
Halobacterium salinarum
Halorutilus salinus
Ca. Nanohalovita haloferacivicina
Ca. Nanohalobium constans
Ca. Nanohalococcus occultus
Methanonatronarchaeum thermophilum
Methanosarcina barkeri MS
Methanohalophilus mahii DSM 5219
Ca. Methanoperedens nitroreducens
Methanothrix thermoacetophila PT
Methanocella conradii HZ254
Methermicoccus shengliensis DSM 18856
Methanosphaerula palustris E1-9c
Methanoregula boonei 6A8
Methanospirillum purgamenti
Methanoplanus endosymbiosus
Methanovulcanius yangii
Methanocalculus alkaliphilus
Methanocorpusculum labreanum Z
Ferroglobus placidus DSM 10642
Archaeoglobus fulgidus DSM 4304
Ca. Wukongarchaeota archaeon
Ca. Heimdallarchaeum aukensis
Ca. Jordarchaeaceae archaeon
Ca. Freyarchaeota archaeon SRVP19_Freyarchaeia-1
Ca. Jordarchaeales archaeon
consensus>70

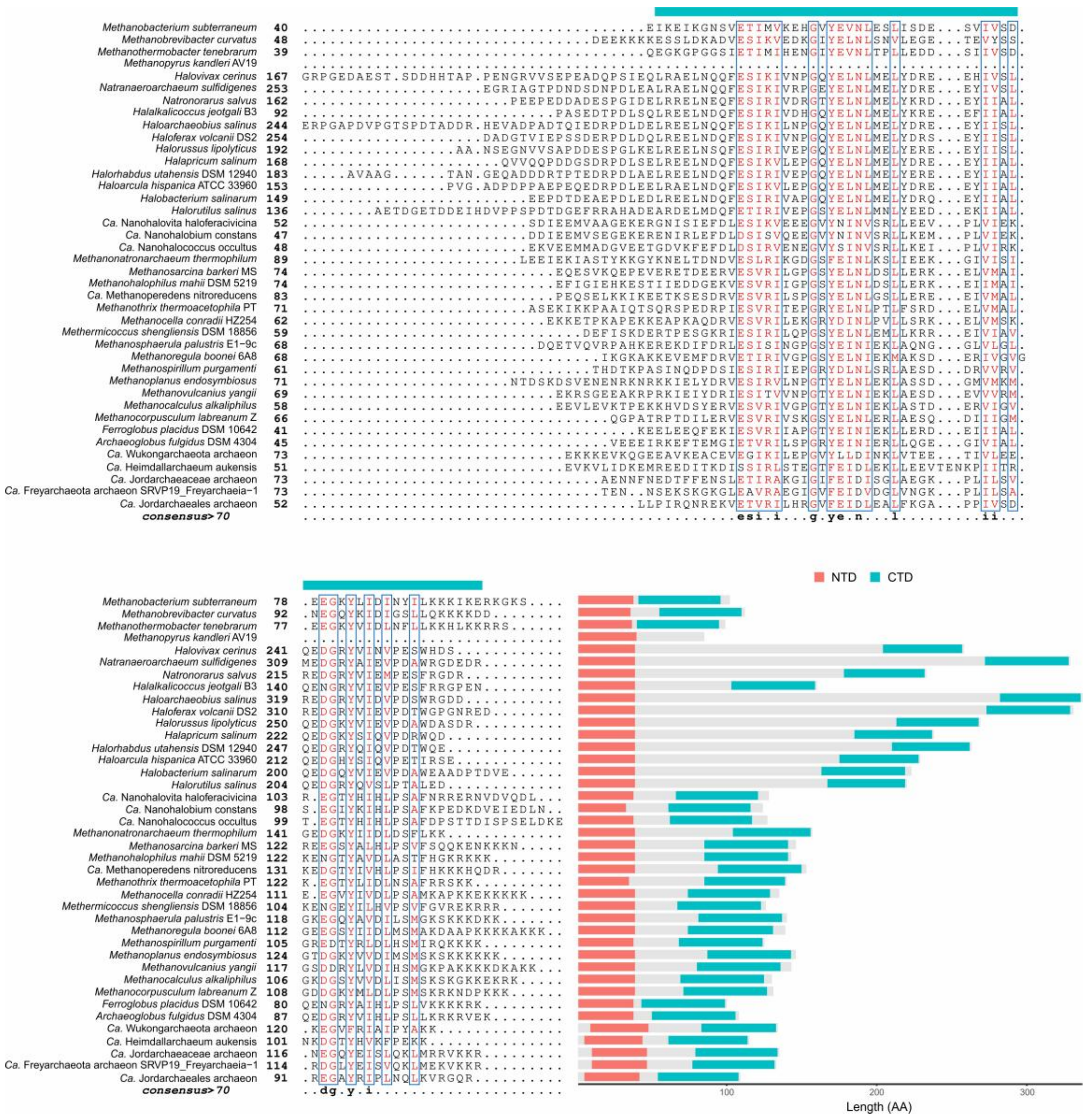

**Supplementary Figure 3: Multiple sequence alignment of DipC.**

40 sequences were picked from the species present in Fig. 1b. Sequences were aligned by Clustal Omega (v1.2.4)[6] and visualized by ESPript (v3.0)[7] with default parameters. The N terminal domain (NTD, 10-50 AA of alignment) and C terminal domain (CTD, 360-420 AA of alignment) are labelled by red and blue bar respectively. A schematic graph to show the length of DipC is put after the alignment, the original positions of NTD and CTD in the proteins are also exhibited.

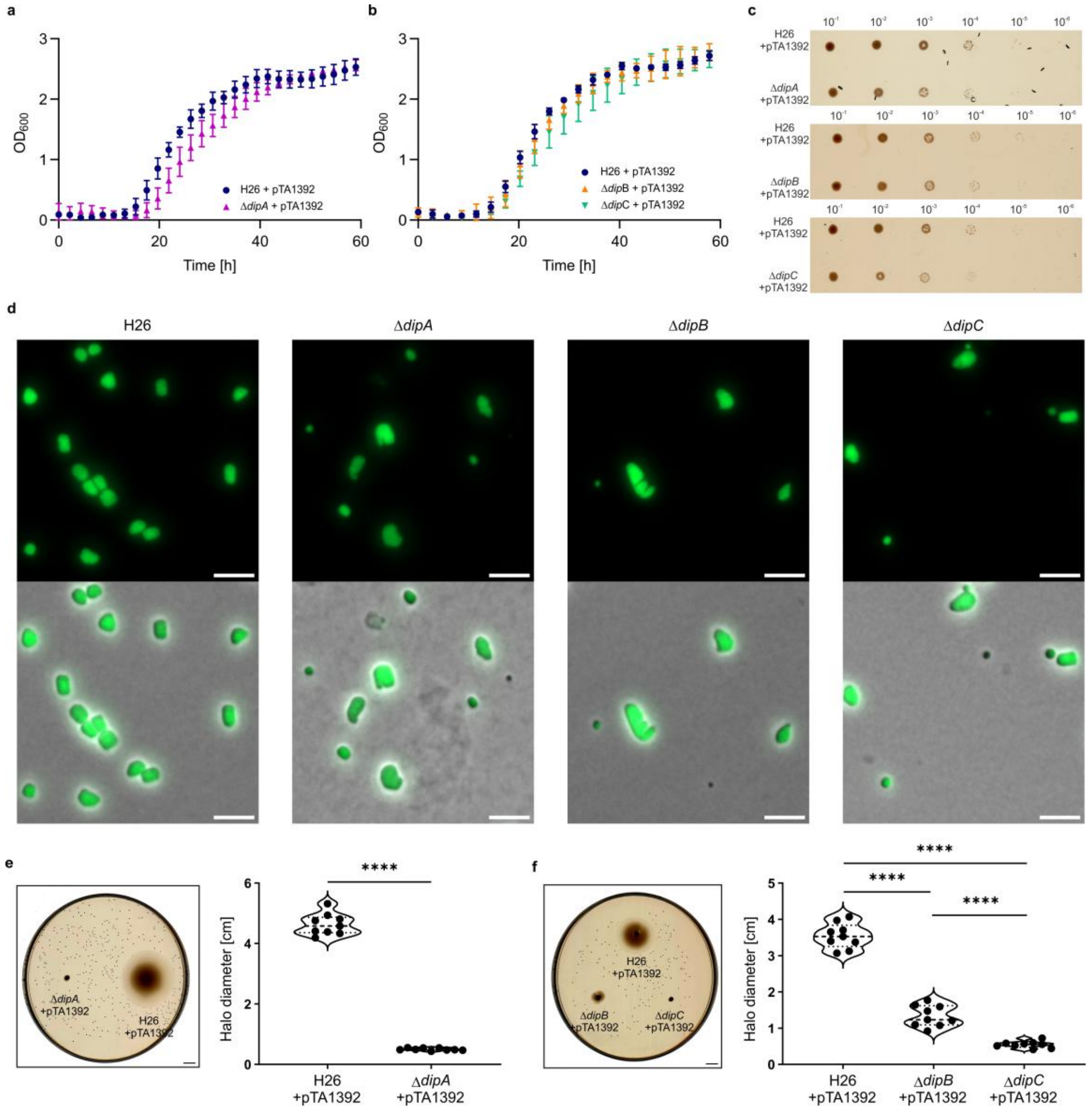

**Supplementary Figure 4: Phenotypes of Δ*dipA*, Δ*dipB* and Δ*dipC*.**

**a-b**, Growth curve of *H. volcanii* wild type H26 and deletion mutants of *dipA*, *dipB* and *dipC* in liquid Hv-Cab medium. Mean of 3 independent replicates are plotted. Error bars, standard deviation (SD). **c**, Spot dilution assay of H26, Δ*dipA*, Δ*dipB* and Δ*dipC* strains. Based on the starting $OD_{600}$ of 0.2, serial dilution of cultures until $10^{-6}$ were made. The diluted cells were spotted on Hv-Ca plates. **d**, Fluorescence microscopy of H26, Δ*dipA*, Δ*dipB* and Δ*dipC* cells at exponential phase ($OD_{600}$ about 0.1) in Hv-Cab medium. 50 µg/mL uracil was added to complement the uracil auxotrophy. Before imaging, cell DNA were stained with 5 µM nucleic acid dye SYTO 13 Green (Thermo Fisher Scientific) for 10 min. Scale bar, 5 µm. **e-f**, Motility assays on semi-solid Hv-Ca plates of different *H. volcanii* strains. For each strain, 3 technical and 3 biological replicates were performed. In violin plot, the 25$^{th}$ and 75$^{th}$ percentile is indicated by dotted line, the median is indicated as dashed line. Significance was tested by two-tailed t-test. **** $P \leq 0.0001$, n = 9. Scale bar, 1 cm.

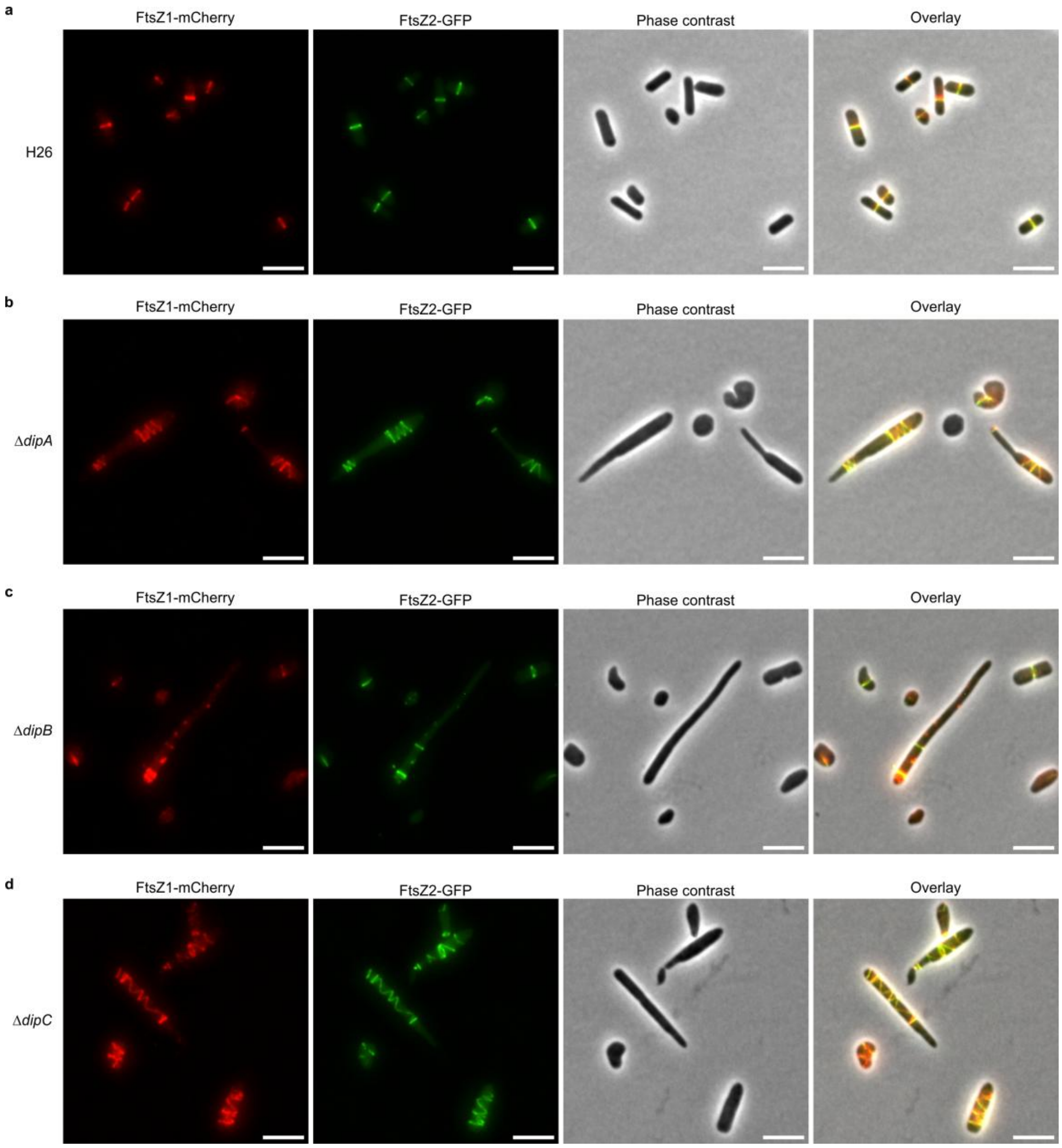

**Supplementary Figure 5: FtsZ1 and FtsZ2 dual localization in wild-type and knockout strains.**

Microscopy of FtsZ1-mCherry + FtsZ2-GFP co-localization (pIDJL134) in (**a**) H26, (**b**) Δ*dipA*, (**c**) Δ*dipB* and (**d**) Δ*dipC* cells. All strains were grown in liquid Hv-Cab medium + 0.25 mM tryptophan. Cells were imaged at exponential phase ($OD_{600}$ about 0.1). All experiments were performed in 3 independent replicates. Scale bars, 5 µm.

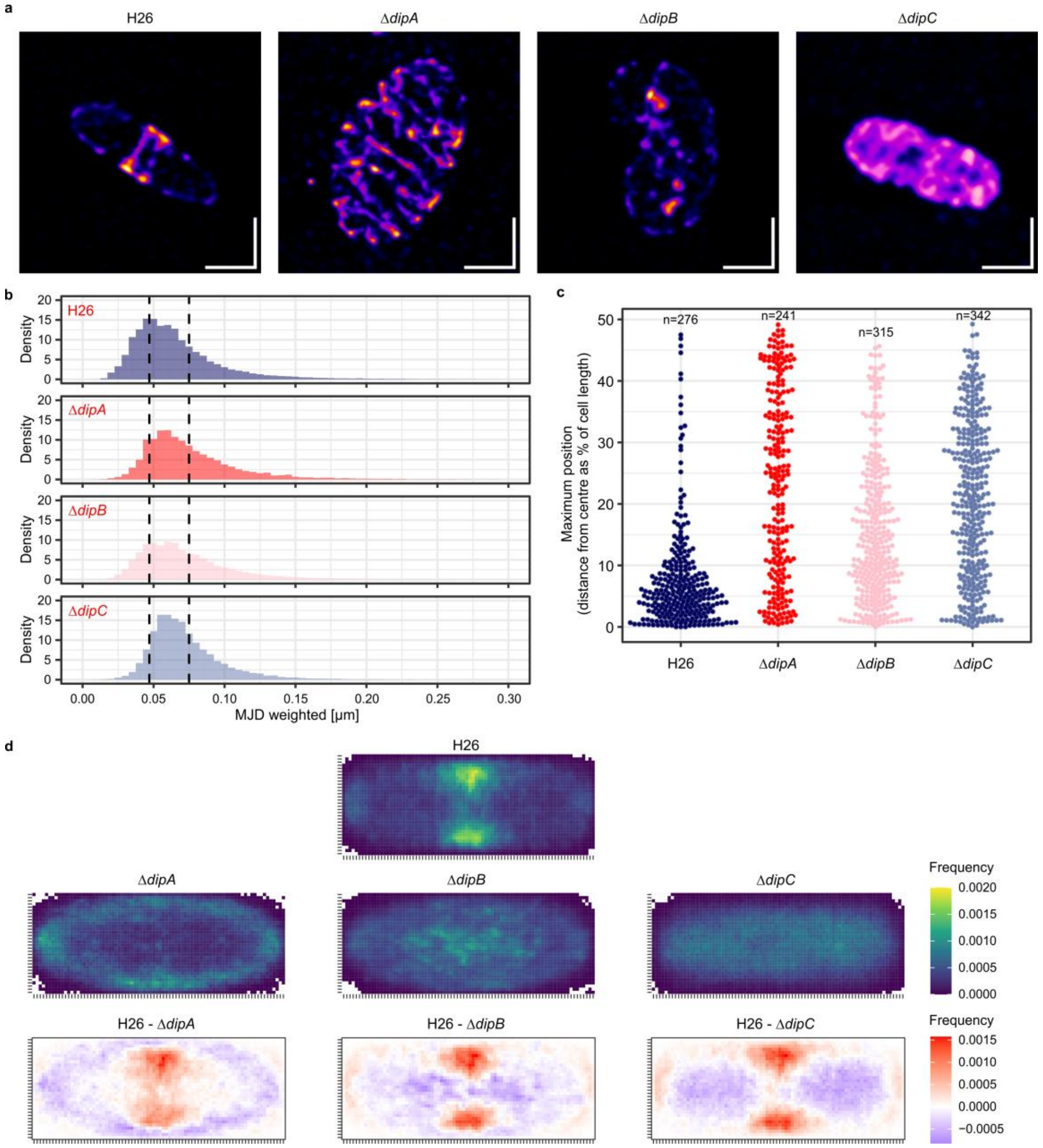

**Supplementary Figure 6: 3D-structured illumination microscopy and single particle tracking of FtsZ1 in H26, Δ*dipA*, Δ*dipB* and Δ*dipC* strains.**

**a,** Representative cells visualized as maximum projection of all 3D-SIM image slices showing different localizations of FtsZ1-HaloTag dyed with R110 in *H. volcanii* H26, Δ*dipA*, Δ*dipB*, and Δ*dipC*. Cells were grown in Hv-Cab medium to an $OD_{600}$ of approximately 0.1, and induced overnight with 0.2 mM tryptophan. Scale bars indicate 1 µm. **b**, Histograms comparing the distribution of weighted mean jump distance (MJD) of all FtsZ1-HaloTag tracks between different strains. The black dotted line indicates population peaks with reference to H26 wild type. **c**, Distribution of the position of the cell segment exhibiting the highest density of FtsZ1-HaloTag tracks along the long axis of the cell. Distance is measured relative to mid-cell (0). Each plotted point represents one cell. **d**, Localization heatmaps showing distribution of all segments in tracks in the average cell obtained by compressing all cells into uniform dimensions in all strains (less tracks in blue to more tracks in yellow). Further, differences are highlighted between the deletion strains and H26 by subtracting the average cells from each other. Red indicates higher probability of presence in H26, purple indicates higher probability of presence in mutants.

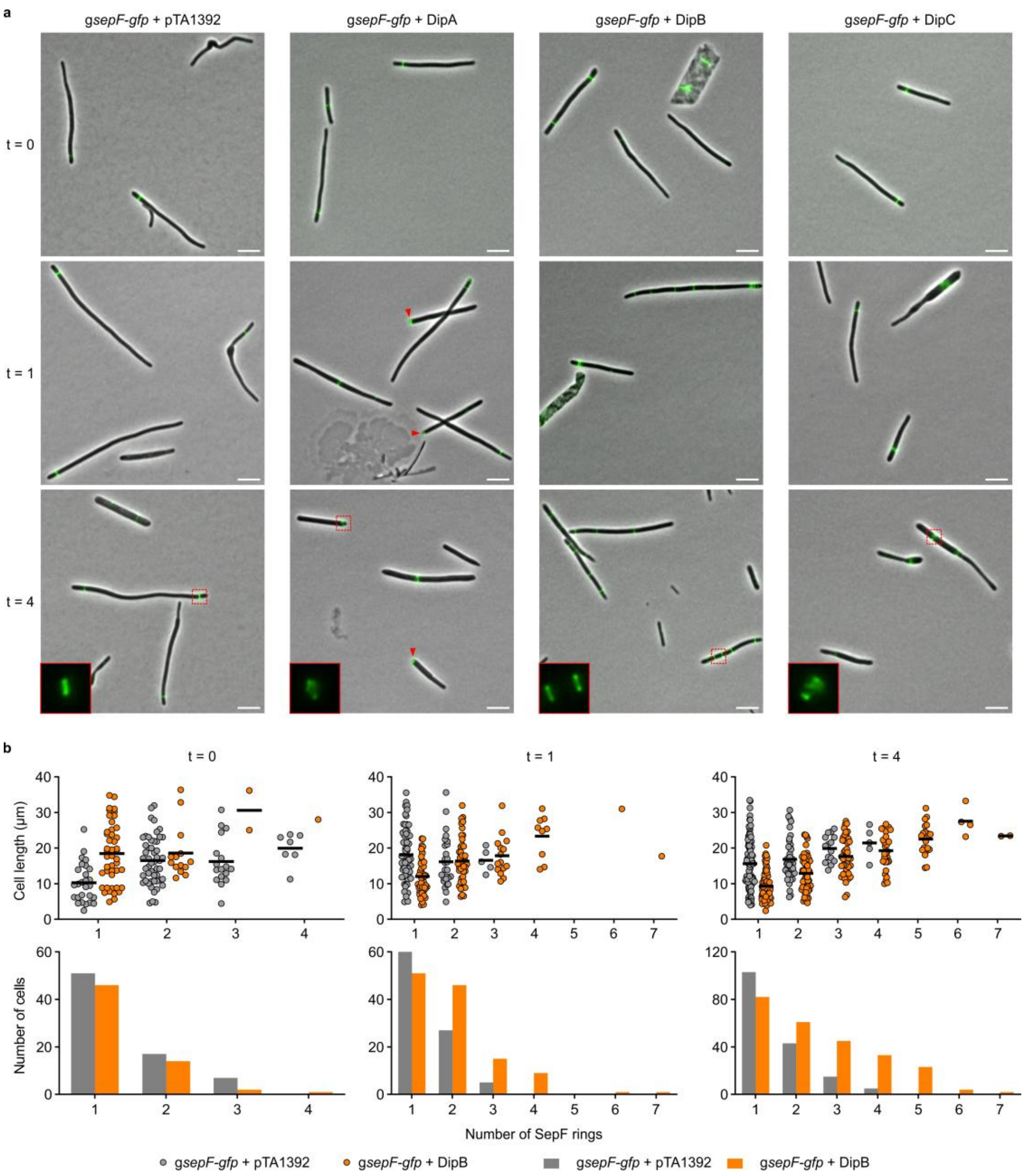

**Supplementary Figure 7: Effect of Dip proteins overexpression on SepF localization.**

**a**, Fluorescence microscopy of *H. volcanii* HTQ959 (*gsepF-gfp*, with genomic integration of a *gfp* tag to *sepF* C-terminal) cells with empty vector (pTA1392) and Dip protein overexpression plasmids (DipA, pSVA6849; DipB, pSVA6848; DipC, pSVA6847). Cells were imaged before induction (t = 0) and different time points (1 and 4 h) after induction. The expression of Dip proteins was induced by adding 1 mM tryptophan. Fluorescent channel image of cells in dashed boxes are magnified at lower left. Representative polar foci are indicated by red arrowheads. Scale bars, 5 µm. **b**, Upper row: cell length in relation to the number of SepF-GFP rings assembled per cell. Mean values are shown as dark lines. Lower row: distribution of SepF rings among the analyzed cells at the indicated time points. The plots summarize the results from three independent experiments per strain and per time point.

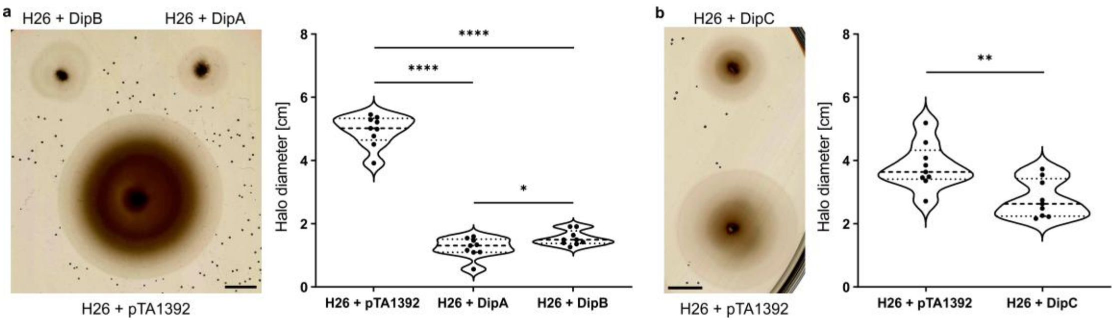

**Supplementary Figure 8: Effect of Dip proteins overexpression on cell motility.**

Motility assays on semi-solid Hv-Ca + 1 mM tryptophan plates of (**a**) DipA, DipB and (**b**) DipC overexpression in H26 cells. For each strain, 3 technical and 3 biological replicates were performed. In violin plot, the $25^{th}$ and $75^{th}$ percentile is indicated by dotted line, the median is indicated as dashed line. Significance was tested by two-tailed t-test. * $P \leq 0.05$, ** $P \leq 0.01$, **** $P \leq 0.0001$, $n = 9$. Scale bar, 1 cm.

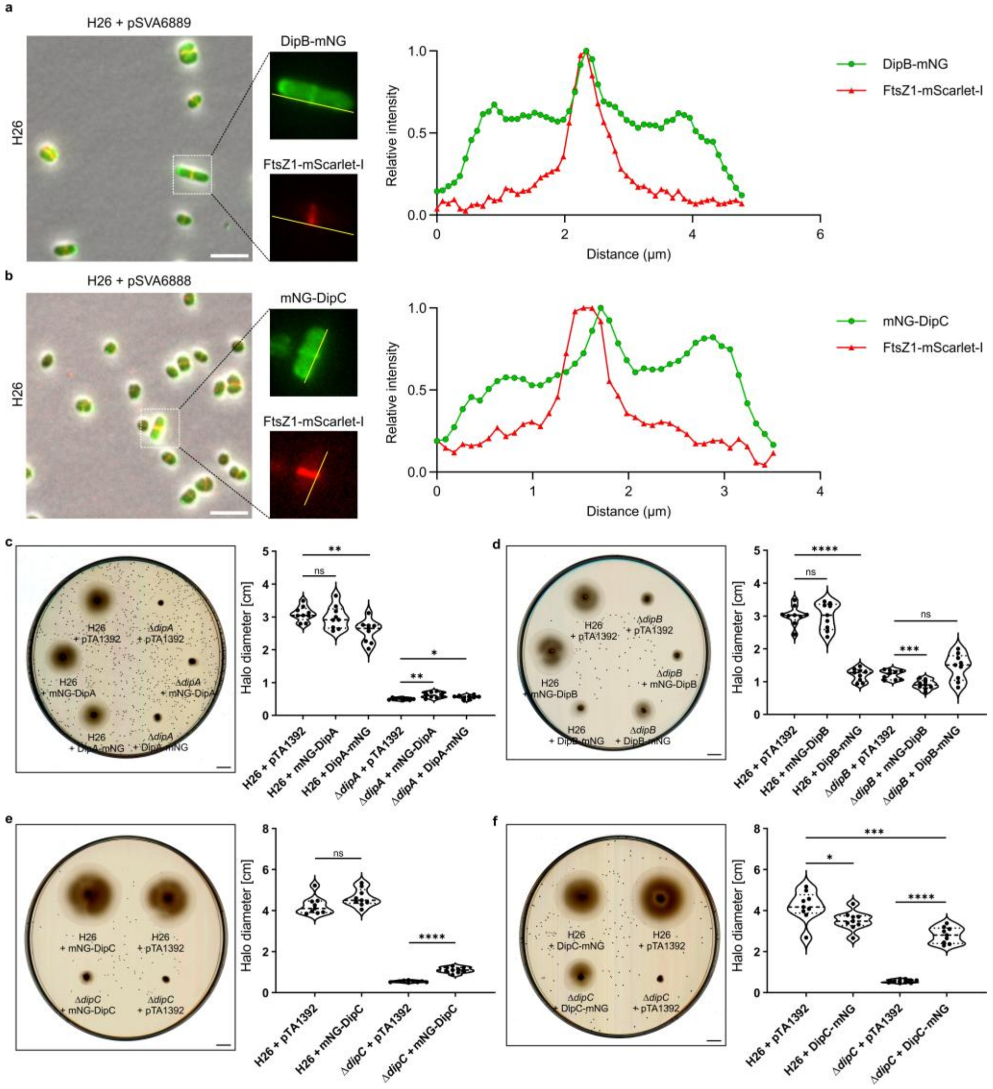

**Supplementary Figure 9: Co-localization of DipB/C with FtsZ1 and motility effect of mNG fused Dip proteins on H26 and KO strains.**

Microscopy of (**a**) DipB-mNG + FtsZ1-mScarlet-I co-localization (pSVA6889) and (**b**) mNG-DipC + FtsZ1-mScarlet-I co-localization (pSVA6888) in H26. All strains were grown in liquid Hv-Cab medium + 0.25 mM tryptophan, and another 0.75 mM tryptophan was added 1 h before imaging. Cells were imaged at exponential phase ($OD_{600}$ about 0.1). The relative signal intensity of mNG and mScarlet along a linear region of interest (ROI) of the representative cell are plotted. The representative cell is highlight by dashed boxes, and the ROI is indicated by a yellow line. Scale bars, 5 µm. **c-f**, Motility assays on semi-solid Hv-Ca + 1 mM tryptophan plates of indicated strains with empty plasmid or mNG fused Dip protein expression plasmids (mNG-DipA, pSVA6825; DipA-mNG, pSVA6826; mNG-DipB, pSVA6823; DipB-mNG, pSVA6824; mNG-DipC, pSVA6455; DipC-mNG, pSVA13781). For each strain, 3 technical and 3 biological replicates were performed. In violin plot, the 25$^{th}$ and 75$^{th}$ percentile is indicated by dotted line, the median is indicated as dashed line. Significance was tested by two-tailed t-test. * $P \leq 0.05$, ** $P \leq 0.01$, *** $P \leq 0.001$, **** $P \leq 0.0001$, ns $P > 0.05$, n = 9. Scale bar, 1 cm.

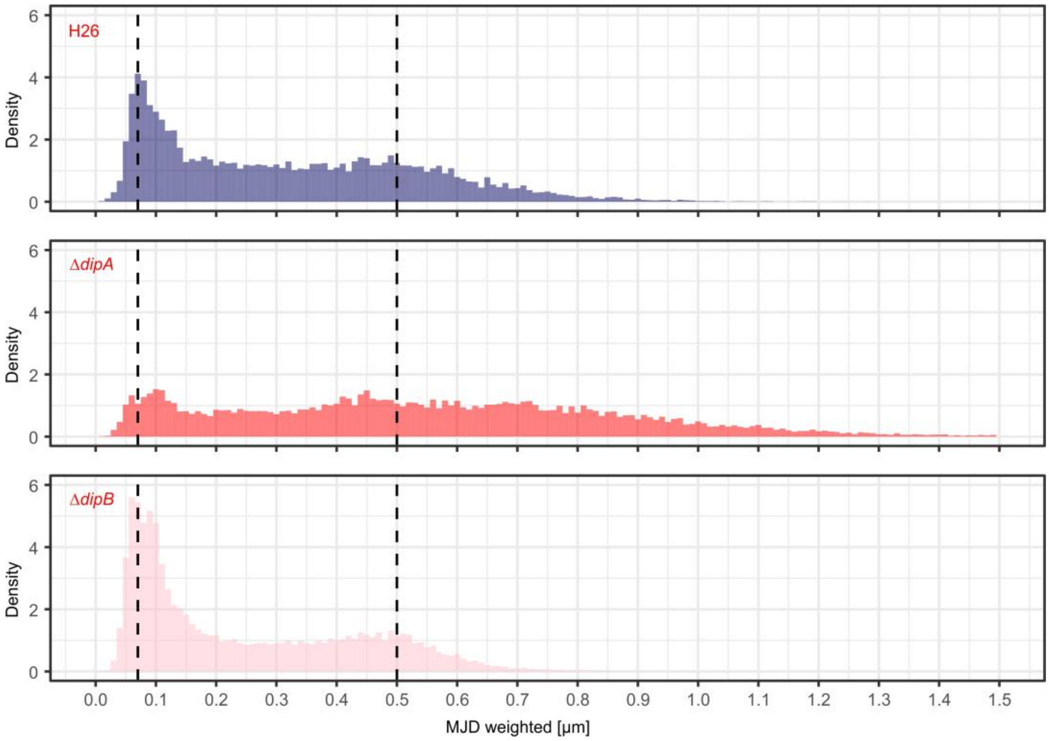

**Supplementary Figure 10: Single particle tracking of DipC in H26, Δ*dipA* and Δ*dipB* strains.**

Histograms comparing the distribution of weighted MJD of all HaloTag-DipC tracks between the different strains. The black dotted line indicates population peaks with reference to H26 wild type. Cells were grown in Hv-Cab medium to an $OD_{600}$ of approximately 0.1, and induced overnight with 0.2 mM tryptophan.

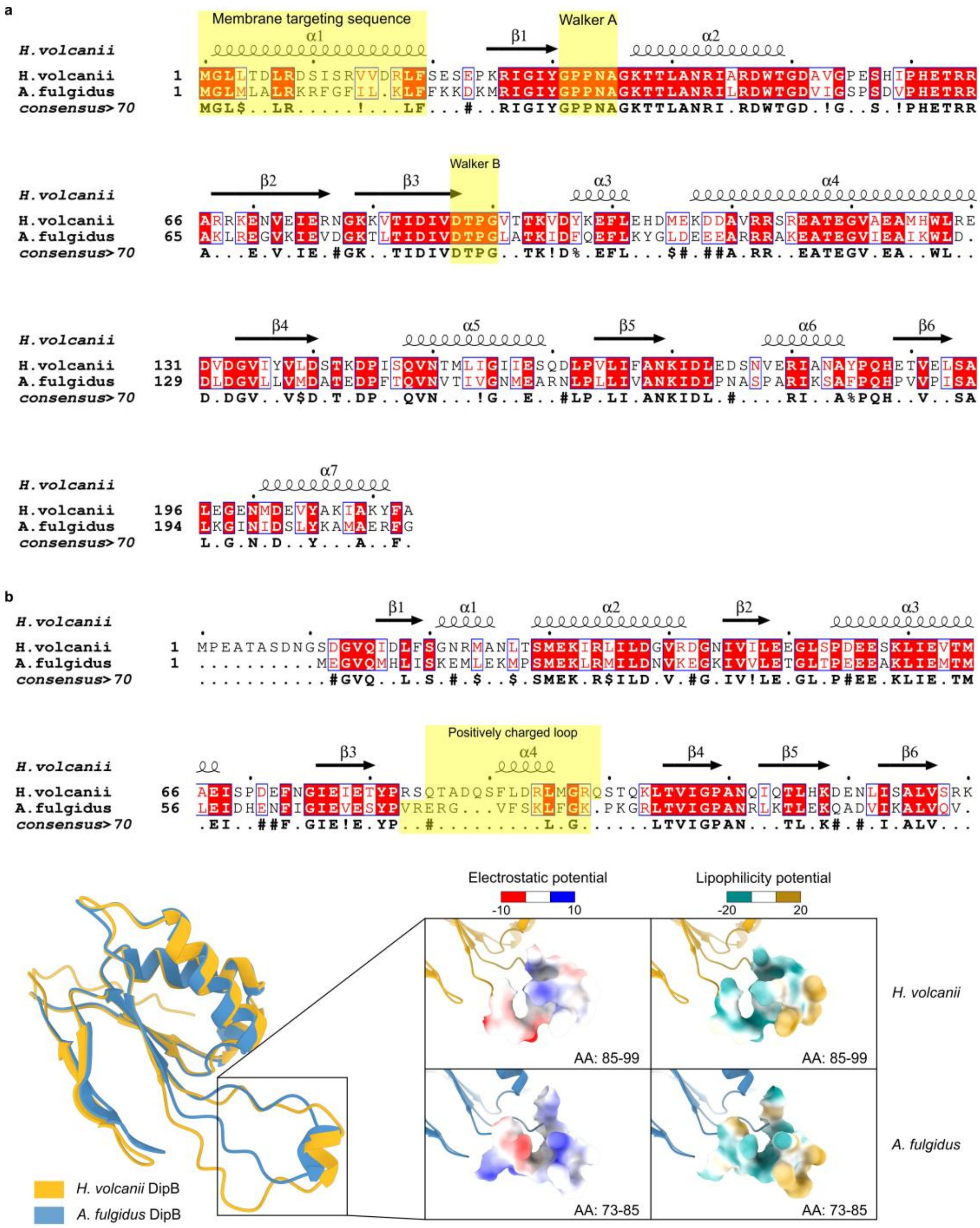

**Supplementary Figure 11: Sequence alignments of DipA and DipB from *H. volcanii* and *A. fulgidus*.**

Sequence alignments of (**a**) DipA and (**b**) DipB from *H. volcanii* and *A. fulgidus*. Sequences were aligned by Clustal Omega (v1.2.4)[6] and visualized by ESPript (v3.0)[7] with default parameters. Secondary structure elements were annotated using 3D structures of *H. volcanii* DipA and DipB, which were predicted by AlphaFold 3[2]. Membrane targeting sequence, Walker A and Walker B motif of DipA, and positively charged loop of DipB are highlight in yellow box. Structures of *H. volcanii* and *A. fulgidus* DipB predicted with AlphaFold 3[2] were aligned by UCSF ChimeraX[3] matchmaker tool with Needleman-Wunsch alignment algorithm and default parameters.

The root mean square deviation (RMSD) between 99 pruned atom pairs is 0.773 Å (across all 115 pairs: 1.972 Å). The loop region of DipB is magnified, the surface is colored according to the electrostatic potential and lipophilicity potential by UCSF ChimeraX[3].

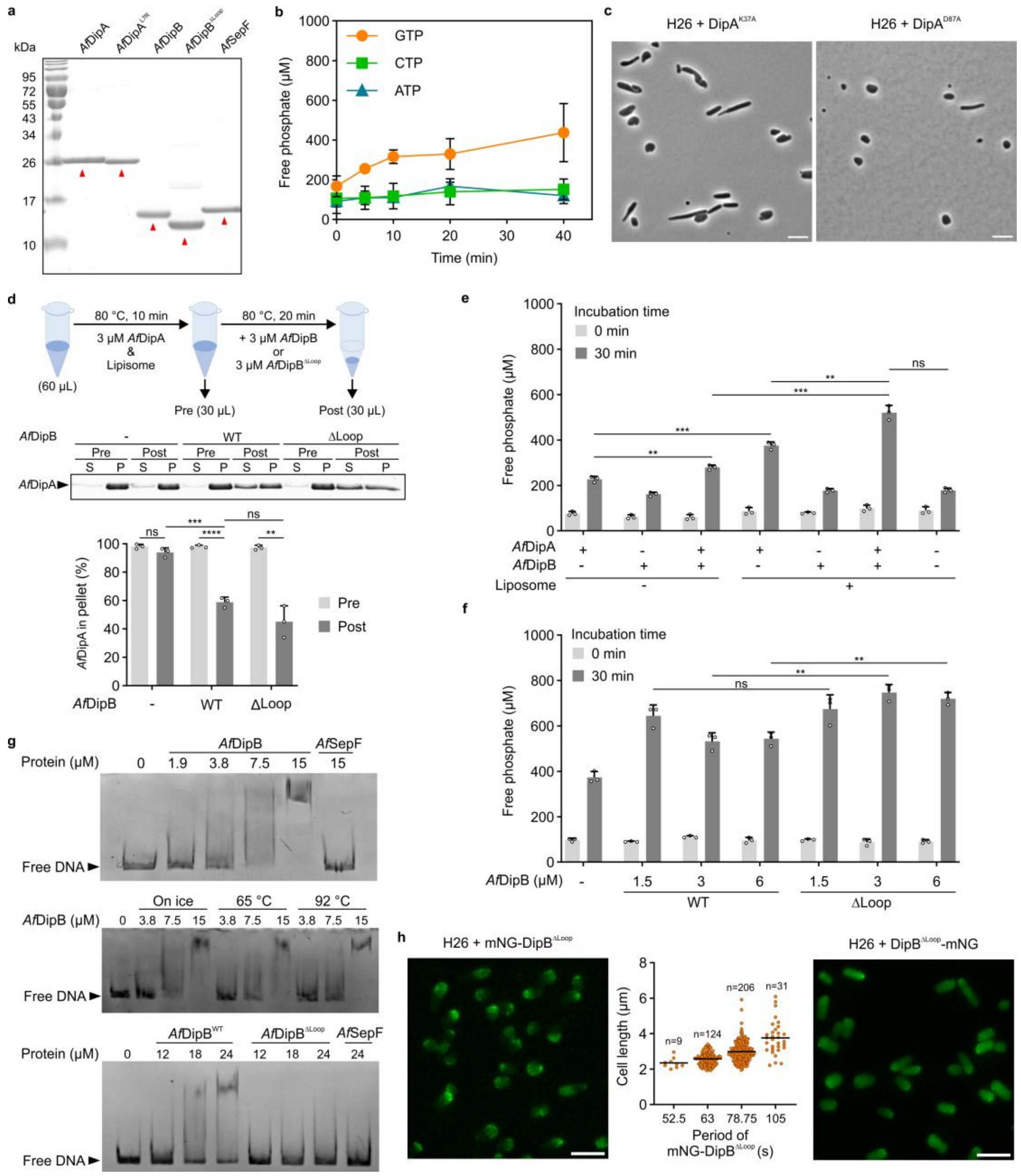

**Supplementary Figure 12: GTPase activity of DipA and DNA binding of DipB.**

**a**, Representative 15% SDS-PAGE gel for purified proteins. The desired bands for each protein are indicated by red triangles. **b**, Hydrolysis of GTP, CTP; and ATP by *Af*DipA. Purified protein and nucleoid were inoculated for the indicated time at 70 °C in a 25 µL reaction mixture. Free phosphate concentration was quantified using the malachite green-ammonium molybdate colorimetric assay (mean ± SD, n = 3). **c**, Phase-contrast images of *H. volcanii* wild-type H26 cells expressing DipA$^{K37A}$ and DipA$^{D87A}$ in liquid Hv-Cab medium + 1 mM tryptophan. **d**, Effect of *Af*DipB on *Af*DipA membrane binding. 3 µM *Af*DipA was first inoculated with liposomes (0.5 mg/mL) at 80 °C in a 60 µL reaction mixture. After 10 min, 3 µM *Af*DipB or *Af*DipB$^{\Delta Loop}$ was added to half of the reaction mixture and incubated for another 20 min. Sedimentation assay was did in the same way as Fig. 5f. Upper panel: Schematic diagram of the assy. Middle panel: Representative 15% SDS-PAGE gel for the liposome-sedimentation of *Af*DipA. Lower panel: Plot for the percentage of pellet fraction of *Af*DipA (mean ± SD, n = 3). **e**, Free phosphate concentration of the reaction mixture. Experiment setting was the same as Fig. 5i. Free phosphate concentration was quantified (mean ± SD, n = 3). **f**, Free phosphate concentration of the reaction mixture. Experiment setting was the same as Fig. 5j. Free phosphate concentration was quantified (mean ± SD, n = 3). **g**, Electrophoretic mobility shift assays (EMSAs) of dsDNA fragments incubated with *Af*DipB, *Af*DipB$^{\Delta Loop}$ or *Af*SepF. **h**, Fluorescence microscopy of *H. volcanii* wild-type H26 cells expressing mNG-DipB$^{\Delta Loop}$ and DipB$^{\Delta Loop}$-mNG. The oscillation periods of mNG-DipB$^{\Delta Loop}$ were plotted, data were taken from 3 independent replicates, mean values are shown as dark lines. Scale bar in (**c**) and (**h**), 5 µm. For (**d**), (**e**), and (**f**), significance was tested by two-tailed t-test. * $P \leq 0.05$, ** $P \leq 0.01$, *** $P \leq 0.001$, **** $P \leq 0.0001$, ns $P > 0.05$, n = 3.

### Supplementary Tables

**Supplementary Table 1: Strains used in this study.**

| Strain name | Background strain | Genotype | Source/ reference |
|---|---|---|---|
| *H. volcanii* | | | |
| H26 | - | *ΔpyrE2* | [8] |
| HTQ909 | H26 | *ΔpyrE2 ΔdipB* | This study |
| HTQ910 | H26 | *ΔpyrE2 ΔdipA* | This study |
| HTQ912 | H26 | *ΔpyrE2 ΔdipC* | This study |
| HTQ971 | H26 | *ΔpyrE2 ftsZ1::gfp* | This study |
| HTQ959 | H26 | *ΔpyrE2 sepF::gfp* | This study |
| *E. coli* | | | |
| DH5α Competent Cells | - | *fhuA2::IS2 Δ(mmuP-mhpD)169 ΔphoA8 glnX44 ϕ80d[ΔlacZ58(M15)] rfbD1 gyrA96 luxS11 recA1 endA1 rphWT thiE1 hsdR17* | New England Biolabs |
| *dam$^{-}$/dcm$^{-}$* Competent Cells | - | *ara-14 leuB6 fhuA31 lacY1 tsx78 glnV44 galK2 galT22 mcrA dcm-6 hisG4 rfbD1 R(zgb210::Tn10)* Tet$^{S}$ *endA1 rspL136 (*Str$^{R}$*) dam13::Tn9 (*Cam$^{R}$*) xylA-5 mtl-1 thi-1 mcrB1 hsdR2* | New England Biolabs |
| NiCo21(DE3) Competent Cells | - | *can::CBD fhuA2 [lon] ompT gal (λ DE3) [dcm] arnA::CBD slyD::CBD glmS6Ala ΔhsdS λ DE3 = λ sBamHIo ΔEcoRI-B int::(lacI::PlacUV5::T7 gene1) i21 Δnin5* | New England Biolabs |
| Rosetta (DE3) | - | F$^{-}$ *ompT hsdS$_{B}$*(r$_{B}^{-}$ m$_{B}^{-}$) *gal dcm* (DE3) pRARE (Cam$^{R}$) | Novagen |

**Supplementary Table 2: Plasmids used in this study.**

| Plasmids | Description | Primers used | Enzymes used | Source/ reference |
|---|---|---|---|---|
| pTA131 | Integrative plasmid with a *pyrE2* selection marker for gene deletions in *H. volcanii* ($Amp^R$) | - | - | [9] |
| pTA1392 | Plasmid for the expression of proteins in *H. volcanii* under control of *p.tnaA* and *pyrE2*, *hdrB* selection markers ($Amp^R$) | - | - | [10] |
| pSVA5910 | Plasmid for expression of FtsZ1-GFP under the control of its native promotor ($Amp^R$) | - | - | [1] |
| pSVA5956 | Plasmid for expression of FtsZ2-GFP under the control of its native promotor with a *pyrE2* selection marker ($Amp^R$) | - | - | [1] |
| pSVA13504 | Plasmid for expression of SepF-GFP under the control of its native promotor with a *pyrE2* selection marker ($Amp^R$) | - | - | [1] |
| pSVA13564 | Plasmid for expression of CdpB1-mNeongreen under the control of its native promotor ($Amp^R$) | - | - | [11] |
| pSVA13565 | Plasmid for expression of CdpB2-mNeongreen under the control of its native promotor ($Amp^R$) | - | - | [11] |
| pSVA13785 | Integrative plasmid for the generation of a *dipC* deletion strain ($Amp^R$) | 13820, 13821, 13822, 13823 | BamHI, in vivo ligation | This study |
| pSVA6813 | Integrative plasmid for the generation of a *dipB* deletion strain ($Amp^R$) | 14266, 14267, 14268, 14269 | In vivo ligation | This study |
| pSVA6814 | Integrative plasmid for the generation of a *dipA* deletion strain ($Amp^R$) | 14270, 14271, 14272, 14273 | In vivo ligation | This study |
| pSVA6847 | Plasmid for expression of DipC | 13812, | NdeI, | This study |

| | | | | |
|---|---|---|---|---|
| | under the control of a tryptophan inducible promotor (Amp$^R$) | 14892 | BamHI | |
| pSVA6848 | Plasmid for expression of DipB under the control of a tryptophan inducible promotor (Amp$^R$) | 14294, 14893 | NdeI, BamHI | This study |
| pSVA6849 | Plasmid for expression of DipA under the control of a tryptophan inducible promotor (Amp$^R$) | 14298, 14894 | NdeI, BamHI | This study |
| pSVA5997 | Plasmid for the expression of proteins with a C-terminal mNeonGreen-tag and a *pyrE2*, *hdrB* selection markers (Amp$^R$) | - | - | [11] |
| pSVA5998 | Plasmid for the expression of proteins with a N-terminal mNeonGreen-tag and a *pyrE2*, *hdrB* selection markers (Amp$^R$) | - | - | [11] |
| pSVA6455 | Plasmid for expression of mNeonGreen-DipC under the control of a tryptophan inducible promotor (Amp$^R$) | 13444, 13445 | NheI, BamHI | This study |
| pSVA13781 | Plasmid for expression of DipC-mNeonGreen under the control of a tryptophan inducible promotor (Amp$^R$) | 13812, 13813 | NdeI, BamHI | This study |
| pSVA6823 | Plasmid for expression of mNeonGreen-DipB under the control of a tryptophan inducible promotor (Amp$^R$) | 14292, 14293 | NheI, BamHI | This study |
| pSVA6824 | Plasmid for expression of DipB-mNeonGreen under the control of a tryptophan inducible promotor (Amp$^R$) | 14294, 14295 | NdeI, BamHI | This study |
| pSVA6825 | Plasmid for expression of mNeonGreen-DipA under the control of a tryptophan inducible promotor (Amp$^R$) | 14296, 14859 | NheI, BamHI | This study |
| pSVA6826 | Plasmid for expression of DipA-mNeonGreen under the control of a tryptophan inducible promotor (Amp$^R$) | 14298, 14299 | NdeI, BamHI | This study |
| pSVA6845 | Plasmid for expression of DipA$^{K37A}$-mNeongreen under | 14888, 14889 | In vitro ligation | This study |

| | | | | |
|---|---|---|---|---|
| | the control of a tryptophan inducible promotor ($Amp^R$) | | | |
| pSVA6846 | Plasmid for expression of $DipA^{D87A}$-mNeongreen under the control of a tryptophan inducible promotor ($Amp^R$) | 14890, 14891 | In vitro ligation | This study |
| pSVA6850 | Plasmid for expression of $DipA^{K37A}$under the control of a tryptophan inducible promotor ($Amp^R$) | 14888, 14889 | In vitro ligation | This study |
| pSVA6851 | Plasmid for expression of $DipA^{D87A}$ under the control of a tryptophan inducible promotor ($Amp^R$) | 14890, 14891 | In vitro ligation | This study |
| pIDJL134 | Plasmid for dual expression of FtsZ2-GFP and FtsZ1-mCherry under the control of a tryptophan inducible promotor ($Amp^R$) | - | - | [12] |
| pSVA3943 | Plasmid to express two proteins, one with a C-terminal GFP tag and one with a C-terminal mCherry tag under the control of a tryptophan inducible promotor, based on pTA1392 ($Amp^R$) | - | - | [1] |
| pSVA3944 | Plasmid to express two proteins, one with a N-terminal GFP tag and one with a C-terminal mCherry tag under the control of a tryptophan inducible promotor, based on pTA1392 ($Amp^R$) | 6597, 8061 (GFP); 8062, 8063 (mCHerry) | NdeI, PciI (GFP); BamHI, NotI (mCHerry) | This study |
| pSVA6881 | Plasmid to express two proteins, one with a C-terminal mNeonGreen tag and one with a C-terminal mScarlet-I tag under the control of a tryptophan inducible promotor, based on pSVA3943 ($Amp^R$) | 15731, 15732 (mNeonGreen); 15733, 15734 (mScarlet-I) | NheI, EcoRI (mNeonGreen); NsiI, NotI (mScarlet-I) | This study |
| pSVA6882 | Plasmid to express two proteins, one with a N-terminal | 14880, 15716 | NdeI, PciI | This study |

| | | | | |
|---|---|---|---|---|
| | mNeonGreen tag and one with a C-terminal mScarlet-I tag under the control of a tryptophan inducible promotor, based on pSVA3944 ($Amp^R$) | (mNeonGreen); 15733, 15734 (mScarlet-I) | (mNeonGreen); NsiI, NotI (mScarlet-I) | |
| pSVA6888 | Plasmid for dual expression of mNeonGreen-DipC and FtsZ1-mScarlet-I under the control of a tryptophan inducible promotor ($Amp^R$) | 14826, 14827 (DipC); 8092, 8093 (FtsZ1) | PciI, NheI (DipC); EcoRI, BamHI (FtsZ1) | This study |
| pSVA6889 | Plasmid for dual expression of DipB-mNeonGreen- and FtsZ1-mScarlet-I under the control of a tryptophan inducible promotor ($Amp^R$) | 15739, 15740 (DipB); 8092, 8093 (FtsZ1) | NdeI, NheI (DipB); EcoRI, BamHI (FtsZ1) | This study |
| pSVA6897 | Plasmid for expression of mNeongreen-$DipB^{\Delta Loop}$ under the control of a tryptophan inducible promotor ($Amp^R$) | 15755, 15756 | In vitro ligation | This study |
| pSVA6898 | Plasmid for expression of $DipB^{\Delta Loop}$-mNeongreen under the control of a tryptophan inducible promotor ($Amp^R$) | 15755, 15756 | In vitro ligation | This study |
| pTA963 HpaA-halotag | Plasmid for expression of HpaA-HaloTag under the control of a tryptophan inducible promotor ($Amp^R$) | - | - | [13] |
| pHVMB006 | Plasmid for expression of HaloTag-DipC under the control of a tryptophan inducible promotor, based on pSVA6455 ($Amp^R$) | KP2_045, KP2_046; KP2_047, KP2_048 | Gibson assembly | This study |
| pHVMB007 | Plasmid for expression of FtsZ1-HaloTag under the control of a tryptophan inducible promotor, | HP_121, HP_12 | Gibson assembly | This study |

| | | | | |
|---|---|---|---|---|
| | based on pHVMB005(Amp$^{R}$) | 2; HP_123, HP_124 | | |
| pSVA13429 | Plasmid for the expression of proteins with an N-terminal His-SUMO-tag in *E. coli* (Kan$^{R}$) | - | - | [11] |
| pSVA13587 | Plasmid for the expression of His-SUMO-SepF from *A. fulgidus* in *E. coli* (Kan$^{R}$) | - | - | [11] |
| pSVA7402 | Plasmid for the expression of His-SUMO-DipB from *A. fulgidus* in *E. coli* (Kan$^{R}$) | 15807, 15808 | In vivo ligation | This study |
| pSVA7403 | Plasmid for the expression of His-SUMO-DipA from *A. fulgidus* in *E. coli* (Kan$^{R}$) | 15809, 15810 | In vivo ligation | This study |
| pSVA7458 | Plasmid for the expression of His-SUMO-DipB$^{\Delta Loop}$ from *A. fulgidus* in *E. coli* (Kan$^{R}$) | 16266, 16268 | In vitro ligation | This study |
| pSVA7459 | Plasmid for the expression of His-SUMO-DipA$^{L7R}$ from *A. fulgidus* in *E. coli* (Kan$^{R}$) | 16263, 16269 | In vitro ligation | This study |
| pSVA13937 | Integrative plasmid to integrate a genomic *gfp*-tag at the C-terminal of *sepF* (Amp$^{R}$) | 14648, 14649; 14650, 14651 | In vivo ligation | This study |
| pSVA13958 | Integrative plasmid to integrate a genomic *gfp*-tag at the C-terminal of *ftsZ1* (Amp$^{R}$) | 15306, 15307; 15308, 15309 | In vivo ligation | This study |

**Supplementary Table 3: Primers used in this study.**

| Primer | Sequence 5'→3' | Description |
|---|---|---|
| pTA131_fw_seq | GGGGGATGTGCTGCAAGG | Forward primer used for sequencing pTA131/pTA1392 based constructs[14] |
| pTA131_rev_seq | TGACCATGATTACGCCAAGC | Reverse primer used for sequencing pTA131/pTA1392 based constructs[14] |
| 13820 | GGCGAATTGGGTACCAGTTGTCAGCGCTCGAAGGG | Forward primer for the amplification of the up-stream region of *hvo_3012* (*dipC*) with 15 complementary bases to linearized pTA131 |
| 13821 | CGCGGATCCTGGTGGATGTTGGCGCCG | Reverse primer for the amplification of the up-stream region of *hvo_3012* (*dipC*) with BamHI restriction site |
| 13822 | CGCGGATCCGCGCGCTTTCCGCGCCC | Forward primer for the amplification of the down-stream region of *hvo_3012* (*dipC*) with BamHI restriction site |
| 13823 | GGCGGCCGCTCTAGAGCAGGCTGACGGTCGTCTTC | Reverse primer for the amplification of the down-stream region of *hvo_3012* (*dipC*) with 15 complementary bases to linearized pTA131 |
| 14266 | GGCGAATTGGGTACCGTGACGCCGTCGGTCCCGAG | Forward primer for the amplification of the up-stream region of *hvo_3013* (*dipB*) with 15 complementary bases to linearized pTA131 |
| 14267 | CGTTCGTGGGGCCCCGGTGGATCACGCGAAGTAC | Reverse primer for the amplification of the up-stream region of *hvo_3013* (*dipB*) with 15 complementary bases to down-stream region of *hvo_3013* (*dipB*) |
| 14268 | CTTCGCGTGATCCACCGGGGCCCCACGAACGCCCTC | Forward primer for the amplification of the down-stream region of *hvo_3013* (*dipB*) with 15 complementary bases to up-stream region of *hvo_3013* (*dipB*) |
| 14269 | GGCGGCCGCTCTAGAGCGGA | Reverse primer for the |

| | | |
|---|---|---|
| | CGGCCACTCGCCCC | amplification of the down-stream region of *hvo_3013* (*dipB*) with 15 complementary bases to linearized pTA131 |
| 14270 | GGCGAATTGGGTACCTAGCG GTCTCCCCCCACCC | Forward primer for the amplification of the up-stream region of *hvo_3014* (*dipA*) with 15 complementary bases to linearized pTA131 |
| 14271 | CTTCAGGCATGGTGGACCTG GGCCTCCCGCACCTCC | Reverse primer for the amplification of the up-stream region of *hvo_3014* (*dipA*) with 16 complementary bases to down-stream region of *hvo_3013* (*dipB*) |
| 14272 | GTGCGGGAGGCCCAGGTCC ACCATGCCTGAAGCAAC | Forward primer for the amplification of the down-stream region of *hvo_3014* (*dipA*) with 16 complementary bases to up-stream region of *hvo_3013* (*dipB*) |
| 14273 | GGCGGCCGCTCTAGATCCTT CGAACCGTCGGGGAAC | Reverse primer for the amplification of the down-stream region of *hvo_3014* (*dipA*) with 15 complementary bases to linearized pTA131 |
| 13812 | GGAATTCCATATGCCACATCA ATGCACCACGTG | Forward primer for the amplification of *dipC* with a NdeI restriction site |
| 14892 | CGCGGATCCTCAGTCCTCGC GGTTGCCGG | Reverse primer for the amplification of *dipC* with a BamHI restriction site |
| 14294 | CGCCATATGATGCCTGAAGC AACCGCCTC | Forward primer for the amplification of *dipB* with a NdeI restriction site |
| 14893 | CGCGGATCCCTACTTTCGCG AGACGAGCGCGC | Reverse primer for the amplification of *dipB* with a BamHI restriction site |
| 14298 | CGCCATATGATGGGACTGCTC ACAGATTTAAG | Forward primer for the amplification of *dipA* with a NdeI restriction site |
| 14894 | CGCGGATCCTCACGCGAAGT ACTTCGCGATTTTG | Reverse primer for the amplification of *dipA* with a BamHI restriction site |

| | | |
|---|---|---|
| 13444 | CGCGCTAGCCCACATCAATG CACCACGTGCGG | Forward primer for the amplification of *dipC* with a NheI restriction site |
| 13445 | CGCGGATCCTCAGTCCTCGC GGTTGCCGGGG | Reverse primer for the amplification of *dipC* with a BamHI restriction site |
| 13813 | CGCGGATCCGTCCTCGCGGT TGCCGG | Reverse primer for the amplification of *dipC* with a BamHI restriction site |
| 14292 | CGCGCTAGCATGCCTGAAGC AACCGCCTC | Forward primer for the amplification of *dipB* with a NheI restriction site |
| 14293 | CGCGGATCCCTACTTTCGCG AGACGAGC | Reverse primer for the amplification of *dipB* with a BamHI restriction site |
| 14295 | CGCGGATCCCTTTCGCGAGA CGAGCGCGC | Reverse primer for the amplification of *dipB* with a BamHI restriction site |
| 14296 | CGCGCTAGCATGGGACTGCT CACAGATTTAAG | Forward primer for the amplification of *dipA* with a NheI restriction site |
| 14859 | CGCGGATCCTCACGCGAAGT ACTTCGCGATTTTG | Reverse primer for the amplification of *dipA* with a BamHI restriction site |
| 14299 | CGCGGATCCCGCGAAGTACT TCGCGATTTTG | Reverse primer for the amplification of *dipA* with a BamHI restriction site |
| 14888 | CCGGGGCGACGACTCTCGCA AATCGTATC | Forward primer for the generation of the Walker A (K37A) mutation in DipA (on pSVA6826) |
| 14889 | CGTTGGGCGGGCCATATATAC CGATCC | Reverse primer for the generation of the Walker A (K37A) mutation in DipA (on pSVA6826) |
| 14890 | GGGGTTACGACGAAGGTCGA TTACAAGGAG | Forward primer for the generation of the Walker B (D87A) mutation in DipA (on pSVA6826) |
| 14891 | CGGGGTGGCGACGATGTCGA TAGTGAC | Reverse primer for the generation of the Walker B (D87A) mutation in DipA (on pSVA6826) |
| 6597 | GTTCTACATATGAGTAAAGG AGAAGAAC | Forward primer for the amplification of *gfp* with a NdeI restriction site |

| | | |
|---|---|---|
| 8061 | ATCACATGTTTGTATAGTTCATCCATGCCATG | Reverse primer for the amplification of *gfp* with a PciI restriction site |
| 8062 | CGGGATCCCCAATGCATGTGAGCAAGGGCGAGGAGGATAAC | Forward primer for the amplification of *mCherry* with a BamHI and NsiI restriction site |
| 8063 | ATAAGAATGCGGCCGCTTACTTGTACAGCTCGTCCATGCC | Reverse primer for the amplification of *mCherry* with a NotI restriction site |
| 15731 | GCGGCTAGCGTCTCGAAGGGCGAGGAAGAC | Forward primer for the amplification of *mNeonGreen* with a NheI restriction site |
| 15732 | GCGGGAATTCTCACTTGTAGAGCTCGTCCATG | Reverse primer for the amplification of *mNeonGreen* with an EcoRI restriction site |
| 15733 | GCGATGCATGTCTCGAAGGGCGAGGCCGTC | Forward primer for the amplification of *mScarlet-I* with a NsiI restriction site |
| 15734 | GCGGCGGCCGCTCACTTGTAGAGCTCGTCCATGC | Reverse primer for the amplification of *mScarlet-I* with a NotI restriction site |
| 14880 | CGCCATATGGTCTCGAAGGGCGAGGAAGAC | Forward primer for the amplification of *mNeonGreen* with a NdeI restriction site |
| 15716 | GCGACATGTCTTGTAGAGCTCGTCCATGC | Reverse primer for the amplification of *mNeonGreen* with a PciI restriction site |
| 14826 | CGCACATGTATGCCACATCAATGCACCAC | Forward primer for the amplification of *dipC* with a PciI restriction site |
| 14827 | CGCGCTAGCTCAGTCCTCGCGGTTGCCGGG | Reverse primer for the amplification of *dipC* with a NheI restriction site |
| 8092 | CCGGAATTCATGGACTCTATCGTCGGCGAC | Forward primer for the amplification of *FtsZ1* with an EcoRI restriction site |
| 8093 | CGCGGATCCCTCGACGTAGTCGATGTCTTC | Reverse primer for the amplification of *FtsZ1* with a BamHI restriction site |
| 15739 | GCGCATATGATGCCTGAAGCAACCGCCTCCG | Forward primer for the amplification of *dipB* with a NdeI restriction site |

| | | |
|---|---|---|
| 15740 | GCGGCTAGCCTTTCGCGAGACGAGCGCGCTG | Reverse primer for the amplification of *dipB* with a NheI restriction site |
| 15755 | TCGACCCAGAAGCTCACGGTCATC | Forward primer for the generation of the ΔLoop mutation in DipB (on pSVA6823 or pSVA6824) |
| 15766 | CGAACGCGGATACGTCTCGATTTC | Reverse primer for the generation of the ΔLoop mutation in DipB (on pSVA6823 or pSVA6824) |
| 15807 | CGCGAACAGATCGGTGGTATGGAAGGAGTTCAGATGCACTTG | Forward primer for the amplification of *A. fulgidus dipB* (*af_1182)* with 18 complementary bases to linearized pSVA13429 |
| 15808 | GGCCAGTGAATCCGTAACTAAACCTGAACGAGGGCCTTAATC | Reverse primer for the amplification of *A. fulgidus dipB* (*af_1182)* with 17 complementary bases to linearized pSVA13429 |
| 15809 | CGCGAACAGATCGGTGGTATGGGATTGATGCTCGCCCTAAGG | Forward primer for the amplification of *A. fulgidus dipA* (*af_1181)* with 18 complementary bases to linearized pSVA13429 |
| 15810 | GGCCAGTGAATCCGTAATCACCCGAACCTCTCGGCCATTG | Reverse primer for the amplification of *A. fulgidus dipA* (*af_1181)* with 17 complementary bases to linearized pSVA13429 |
| 16266 | CGGATACGACTCAACCTCAATG | Reverse primer for the generation of the ΔLoop mutation in *Af*DipB (on pSVA7402) |
| 16268 | CCGAAAGGGAGGCTGACG | Forward primer for the generation of the ΔLoop mutation in *Af*DipB (on pSVA7402) |
| 16263 | GGCGAGCATCAATCCCATAC | Reverse primer for the generation of the L7R mutation in *Af*DipA (on pSVA7403) |
| 16269 | CGCAGGAAGAGATTCGGATTCATTCTG | Forward primer for the generation of the L7R mutation in *Af*DipA (on pSVA7403) |
| 14648 | GGCGAATTGGGTACCGGCGAGACCTGTCTGAAATC | Forward primer for the amplification of *sepF-gfp* region with 15 complementary bases to linearized pTA131 |
| 14649 | GTGTCGTCCGCCGTCGTTATT | Reverse primer for the |

| | | |
|---|---|---|
| | TGTATAGTTCATCC | amplification of *sepF-gfp* with <u>16 complementary bases</u> to start of down-stream region of *sepF* |
| 14650 | <u>GAACTATACAAATAA</u>CGACG GCGGACGACACCCG | Forward primer for the amplification of the down-stream region of *sepF* with <u>15 complementary bases</u> to end of *gfp* |
| 14651 | <u>GGCGGCCGCTCTAGA</u>GACTG CTCGCCGACCTCCG | Reverse primer for the amplification of the down -stream region of *sepF* with <u>15 complementary bases</u> to linearized pTA131 |
| 15306 | <u>GGCGAATTGGGTACC</u>GGTCA AGGGCATCACCGAAC | Forward primer for the amplification of *ftsZ1-gfp* region with <u>15 complementary bases</u> to linearized pTA131 |
| 15307 | <u>TGCGGGACGGCTCGA</u>TTATTT GTATAGTTCATCC | Reverse primer for the amplification of *ftsZ1-gfp* with <u>15 complementary bases</u> to start of down-stream region of *ftsZ1* |
| 15308 | <u>GAACTATACAAATAA</u>TCGAG CCGTCCCGCACGCAC | Forward primer for the amplification of the down-stream region of *ftsZ1* with <u>15 complementary bases</u> to end of *gfp* |
| 15309 | <u>GGCGGCCGCTCTAGA</u>CCGTC GTCGTCGGCCTCGAC | Reverse primer for the amplification of the down -stream region of *ftsZ1* with <u>15 complementary bases</u> to linearized pTA131 |
| KP2_045 | TGCGCATATGGAAATCGGTA CTGGCTTTC | Forward primer for the amplification of the HaloTag |
| KP2_046 | GTGGGCTAGCGGAAATCTCC AGAGTAGAC | Reverse primer for the amplification of the HaloTag |
| KP2_047 | GGAGATTTCCGCTAGCCCAC ATCAATGC | Forward primer for the linearization of pSVA6455 |
| KP2_048 | TACCGATTTCCATATGCGCAA TAGGTCC | Reverse primer for the linearization of pSVA6455 |
| HP_121 | CTACGTCGAGGGTGGCGGGG GTAGCGGT | Forward primer for the linearization of pTA963 HpaA-halotag |
| HP_122 | TAGAGTCCATGTGGTGGTGG TGGTGGTGC | Reverse primer for the linearization of pTA963 HpaA- |

| | | |
|---|---|---|
| | | halotag |
| HP_123 | CCACCACCACATGGACTCTA TCGTCGGC | Forward primer for the amplification of *ftsZ1* (*hvo_0717*) |
| HP_124 | CCCCGCCACCCTCGACGTAG TCGATGTC | Reverse primer for the amplification of *ftsZ1* (*hvo_0717*) |

### Supplementary Data Legends

**Supplementary Data 1: Archaea genome assembly used in this study.**

The genome assembly information was collected from NCBI database, and the assemblies used in Fig. 1b were filled with orange in cell.

**Supplementary Data 2: The Archaea Dip system found by MacSyFinder.**

The Dip system was searched across the chosen Archaea genomes with MacSyFinder (v2.1.3)[15]. The results were manually inspected, and the proteins expressed from pseudogenes were labeled as red text.

**Supplementary Data 3: The Archaea FtsZ homologs found by hmmsearch.**

The FtsZ1 and FtsZ2 HMM models were build based on previously published data[12], check Methods section for more details. The results were manually inspected, and the proteins expressed from pseudogenes were labeled as red text.

**Supplementary Data 4: The MacSyFinder model of Archaea Dip system.**

The model was created by *msf_data init* in MacSyFinder (v2.1.3)[15] according to the standard protocol of the program.

### Supplementary Movies Legends

**Supplementary Movie 1: Overnight microscopy of wild-type H26, Δ*dipA*, Δ*dipB*, and Δ*dipC* cells in microfluidic chambers.**

H26, Δ*dipA*, Δ*dipB*, and Δ*dipC* cells were transformed with empty plasmid (pTA1392). Cells for each strain were grown in Hv-Cab medium to an $OD_{600}$ about 0.1. Cell culture was then load into a pre-washed microfluidic plate. Images were acquired every 15 min for 16 h at 45 °C with a constant flow of fresh Hv-Cab medium at 5 kPa. Only parts of the complete videos are shown here. All scale bars, 5 µm.

**Supplementary Movie 2: Overnight microscopy of FtsZ1-GFP in wild-type H26, Δ*dipA*, Δ*dipB*, and Δ*dipC* cells in microfluidic chambers.**

H26, Δ*dipA*, Δ*dipB*, and Δ*dipC* cells were transformed with FtsZ1-GFP expression plasmid (pSVA5910). Cells for each strain were grown in Hv-Cab medium to an $OD_{600}$ about 0.1. Cell culture was then load into a pre-washed microfluidic plate. Images were acquired every 15 min for 16 h at 45 °C with a constant flow of fresh Hv-Cab medium at 5 kPa. Only parts of the complete videos are shown here. All scale bars, 5 µm.

**Supplementary Movie 3: 3D-SIM reconstruction images of FtsZ1-GFP in wild-type H26, Δ*dipA*, Δ*dipB*, and Δ*dipC* cells in microfluidic chambers.**

Representative cells visualized as a 3D projection of all 3D-SIM image slices showing different localizations of FtsZ1-HaloTag dyed with R110 in *H. volcanii* H26, Δ*dipA*, Δ*dipB,* and Δ*dipC.* Cells were grown in Hv-Cab medium to an $OD_{600}$ of approximately 0.1, and induced overnight with 0.2 mM tryptophan. All scale bars, 1 µm.

**Supplementary Movie 4: Time-lapse microscopy of mNG tagged Dip proteins in wild-type H26 cells on agarose pad.**

Plasmids expressing mNG-DipA (pSVA6825), DipA-mNG (pSVA6826), mNG-DipB (pSVA6823), DipB-mNG (pSVA6824), mNG-DipC (pSVA6455), DipC-mNG (pSVA13781) were transformed into H26 cells. Cells for each strain were grown in 20 mL Hv-Cab medium to an $OD_{600}$ about 0.1 with 4 h induction of expression by 1 mM tryptophan before imaging. 5 µL cell culture was then load onto the agarose pad. Images were acquired every 15 s for 5 min at 45 °C. For DipA-mNG, images were acquired every 15 s for 10 min at 45 °C. The cells chosen to exhibit in Fig 4e-g were highlight by yellow boxes. All mNG constructs were examined with 3 independent biological replicates. All scale bars, 5 µm.

**Supplementary Movie 5: Overnight microscopy of mNG-DipC in wildtype H26 cells.**

Plasmid expressing mNG-DipB (pSVA6455) was transformed into H26 strain. Cells were grown in Hv-Cab medium to an $OD_{600}$ about 0.1 with 1 h induction of expression by 1 mM tryptophan before imaging. Cell culture was then load into a pre-washed microfluidic plate. Images were acquired every 15 min for 16 h at 45 °C with a constant flow of fresh Hv-Cab medium + 1 mM tryptophan at 5 kPa. Only parts of the complete videos are shown here. All scale bars, 5 µm.

**Supplementary Movie 6: Time-lapse microscopy of mNG tagged Dip proteins in indicated *dip* gene deletion strains on agarose pad.**

Plasmids expressing mNG-DipA (pSVA6825), DipA-mNG (pSVA6826), mNG-DipB (pSVA6823), DipB-mNG (pSVA6824), mNG-DipC (pSVA6455), DipC-mNG (pSVA13781) were transformed into *dip* gene deletion cells. Cells for each strain were grown in 20 mL Hv-Cab medium to an $OD_{600}$ about 0.1 with 4 h induction of expression by 1 mM tryptophan before imaging. 5 µL cell culture was then load onto the agarose pad. Images were acquired every 15 s for 5 min at 45 °C. All mNG constructs were examined with 3 independent biological replicates. All scale bars, 5 µm.

**Supplementary Movie 7: Time-lapse microscopy of DipA$^{K37A}$-mNG and DipA$^{D87A}$-mNG in wildtype H26 cells on agarose pad.**

Plasmids expressing DipA$^{K37A}$-mNG (pSVA6845) and DipA$^{D87A}$-mNG (pSVA6846) were transformed into H26 cells. Cells for each strain were grown in 20 mL Hv-Cab medium to an $OD_{600}$ about 0.1 with 4 h induction of expression by 1 mM tryptophan before imaging. 5 µL cell culture was then load onto the agarose pad. Images were acquired every 15 s for 5 min at 45 °C. All mNG constructs were examined with 3 independent biological replicates. All scale bars, 5 µm.

**Supplementary Movie 8: Time-lapse microscopy of mNG-DipB$^{\Delta Loop}$ and DipB$^{\Delta Loop}$-mNG in wildtype H26 and Δ*dipB* cells.**

Plasmids expressing mNG-DipB$^{\Delta Loop}$ (pSVA6897) and DipB$^{\Delta Loop}$-mNG (pSVA6898) in H26 and Δ*dipB* cells. Cells for each strain were grown in 20 mL Hv-Cab medium to an $OD_{600}$ about 0.1 with 4 h induction of expression by 1 mM tryptophan before imaging. 5 µL cell culture was then load onto the agarose pad. Images were acquired every 15 s for 5 min at 45 °C. All mNG constructs were examined with 3 independent biological replicates. All scale bars, 5 µm

**Supplementary Movie 9: Time-lapse microscopy of mNG tagged Dip proteins in Δ*dipA*, Δ*dipB*, and Δ*dipC* cells on agarose pad.**

Plasmids expressing mNG-DipA (pSVA6825), DipA-mNG (pSVA6826), mNG-DipB (pSVA6823), DipB-mNG (pSVA6824), mNG-DipC (pSVA6455), DipC-mNG (pSVA13781) were transformed into *dip* gene deletion cells. Cells for each strain were grown in 20 mL Hv-Cab medium to an $OD_{600}$ about 0.1 with 4 h induction of expression by 1 mM tryptophan before imaging. 5 µL cell culture was then load onto the agarose pad. Images were acquired every 15 s for 5 min at 45 °C. All mNG constructs were examined with 3 independent biological replicates. All scale bars, 5 µm.

**Supplementary Movie 10: Overnight microscopy of DipB-mNG in wildtype H26 cells.**

Plasmid expressing DipB-mNG (pSVA6824) was transformed into H26 strain. Cells were grown in Hv-Cab medium to an $OD_{600}$ about 0.1 with 1 h induction of expression by 1 mM tryptophan before imaging. Cell culture was then load into a pre-washed microfluidic plate. Images were acquired every 15 min for 16 h at 45 °C with a constant flow of fresh Hv-Cab medium + 1 mM tryptophan at 5 kPa. Only parts of the complete videos are shown here. All scale bars, 5 μm.

.
